# SH2 Domain Protein E sustains perivascular fibroblasts to maintain arterial stability

**DOI:** 10.64898/2026.11.12.766010

**Authors:** Surendra Kumar Anand, Krishna K. Rentachintala, Jaylen E. Moore, Diandra Rufin Florat, Jennifer A. Schumacher, Ivana Zlatanova, Brian L. Black, Saulius Sumanas

## Abstract

Developing arteries must withstand pulsatile blood flow while remaining sufficiently flexible to accommodate changing hemodynamic demands. Endothelial interactions with perivascular cells and the extracellular matrix (ECM) are essential for this balance, but the signaling mechanisms that coordinate them remain incompletely understood. We previously showed that SH2 domain protein E (She), an endothelial adaptor protein implicated in ABL signaling, is required to maintain the size and structural stability of the dorsal aorta (DA) in zebrafish embryos. *she* mutant embryos develop DA collapse and circulatory failure at 4–5 days post-fertilization, yet the underlying cause has not been identified. Here, we investigated the role of She in recruiting ECM-producing perivascular stromal cells to regulate the structural stability of the DA. *She* mutants had fewer collagen-producing perivascular stromal cells between the DA and posterior cardinal vein. This correlated with smaller perivascular *col1a1* and *col5a1* expression domains, reduced *col1a1* transcript and Collagen I pro-peptide levels, and increased DA distensibility. Single-cell RNA sequencing independently identified a reduced relative abundance of a perivascular fibroblast subpopulation. In contrast, DA-associated mural cell number, laminin immunoreactivity, and local *col4a1* transcript levels were not detectably altered. Partial ablation of mural cells alleviated DA narrowing and circulatory failure, while the nitric oxide donor sodium nitroprusside provided partial rescue. These findings support a contribution of mural cell-dependent vascular tone to the collapse of the mechanically vulnerable DA. Mutants also exhibited cardiac chamber enlargement, increased atrial and bulbus arteriosus deformation, atrioventricular regurgitation, and increased Elastin B accumulation in the enlarged bulbus arteriosus. However, endocardial-specific She expression failed to rescue circulatory failure, supporting its requirement in the vascular endothelium. Together, these findings argue that She functions in the vascular endothelium to recruit and/or maintain perivascular stromal cells, which produce fibrillar collagens required for arterial stability. These results reveal a new mechanism by which signals from the vascular endothelium promote ECM support to maintain arterial vascular stability.

---

Keywords: She; arterial stiffness; perivascular fibroblasts; stromal cells; fibrillar collagens; mural cells; dorsal aorta; zebrafish

## Introduction

Developing blood vessels must maintain an open lumen while accommodating pulsatile flow and changes in vascular pressure. This requires coordinated regulation of endothelial behavior, extracellular matrix (ECM) assembly, and interactions with surrounding cells [1–9]. The endothelial layer, interstitial matrix, and basement membrane collectively support the vessel wall and transmit forces between endothelial and perivascular cells [5, 10–18]. Disruption of these interactions can cause abnormal vessel enlargement, excessive compliance, stenosis, or collapse.

The interstitial matrix is composed mainly of fibrillar collagens, including collagen types I and V. These fibrillar collagens co-assemble into heterotypic tensile fibrils that limit excessive tissue deformation and absorb hemodynamic force [18–20]. Perivascular and adventitial fibroblasts are important sources of these collagens [18, 19]. Endothelial signals and local mechanical cues help coordinate the recruitment and function of surrounding mesenchymal cells [21, 22]. However, the role of endothelial cells in maintaining perivascular fibroblast populations and their matrix-producing activity during vascular development remain poorly understood.

Mural cells contribute to vessel maturation by regulating vascular tone and promoting basement membrane assembly [10, 13, 14, 23, 24]. In the zebrafish trunk, mural cells progressively associate with the dorsal aorta (DA) between 2 and 5 days post-fertilization (dpf), coinciding with reduced aortic diameter and increased vessel-wall stabilization [14, 25]. Vessel caliber reflects the balance among intraluminal pressure, wall properties, and forces generated by surrounding cells. A vessel with inadequate structural support may therefore become vulnerable to mural cell-dependent constriction. How endothelial signaling maintains this balance during arterial maturation remains unclear.

Src homology 2 domain-containing E (She) is an intracellular adaptor protein associated with ABL kinase signaling [26]. We previously showed that *she* is specifically expressed in vascular endothelial cells during zebrafish embryogenesis [27]. Loss-of-function studies demonstrated that She is required to maintain vascular lumen size in zebrafish embryos and human endothelial cell culture. Zebrafish *she* mutants initially develop an enlarged DA due to increased proliferation of vascular endothelial cells. SHE depletion similarly enlarges endothelial tubes in three-dimensional culture of human endothelial cells. Despite early enlargement of the DA, blood circulation appears normal in *she* mutants until 4 dpf. At approximately 4-5 dpf, progressive collapse of the DA and cessation of blood flow are observed in *she* mutants [28]. As a result, homozygous *she* mutant embryos died by approximately 7 dpf [28]. However, the cause of this later phenotype remains unclear.

Here, we investigated how loss of She affects the stability of the developing DA. We show that increased pulsatile diameter change precedes overt circulatory failure and is accompanied by reduced perivascular stromal cell abundance and fibrillar collagen gene expression. Although mural cell abundance is unchanged, partial mural cell ablation prevents DA collapse and circulation failure, and pharmacological vasodilation provides partial rescue. We also examine cardiac, outflow tract, and pronephric abnormalities and test whether endocardial She expression or reduction of Elastin B can rescue the *she* mutant phenotype. These findings support a model in which endothelial She enables the arterial wall to resist mural cell-dependent constrictive forces during vascular maturation by regulating perivascular stromal cell number and modulating perivascular collagen deposition.

## Materials and Methods

### Ethics statement

All zebrafish husbandry and experimental procedures were approved by the Institutional Animal Care and Use Committee (IACUC).

### Zebrafish lines

Previously generated *she^ci26^* and *Tg(fli1:she-2A-mCherry)^ci34^* lines were used in this study [28]. Additional lines included *Tg(kdrl:GFP)^s843^* [29], *Tg(kdrl:Has.HRAS-mCherry)^s896^* [30], *Ki(pdgfrb:gal4)^ion114^* [31], *Tg(UAS:mCherry-NTR)^c264^* [32], *Tg(fli1a:lifeact-GFP)^mu240^* [33], *Tg(fli1:nGFP)^y7^* [34], *Tg(tjp1a:eGFP)^pd1252^* [35], and *Tg(pdgfrb:GFP)^ncv22^* [36]. Embryos were maintained under standard conditions. Where indicated, embryos were treated with 0.003% 1-phenyl-2-thiourea (PTU) beginning at 24 hpf to inhibit pigmentation.

### Generation of the Tg(endo83:she-2A-mCherry) line

We generated a transgenic zebrafish line using the endocardial-specific *endo83* enhancer (detailed characterization of the enhancer will be published elsewhere). The *endo83:she-2A-mCherry* construct was generated using the Gateway recombination system (Invitrogen). The endo83 enhancer was amplified by PCR and subcloned into the P4P1R 5’ entry vector by BP recombination. Subsequently, the previously generated middle-entry vector *pDONR221-she* [28] was recombined by an LR reaction with endo83-P4P1R and p3E-2A-mcherrypA into the pDestTol2pA2-alpha-crystallin destination vector [37]. Embryos in the *she^ci26^* mutant background were injected with 20 ng of *endo83:she-2A-mCherry* DNA and 150 ng of Tol2 mRNA (Tol2-mediated transgenesis). Transgenic embryos were identified by endocardial mCherry expression and raised to establish stable lines.

### Brightfield and confocal imaging

Live and fixed embryos were imaged using a Nikon AX confocal microscope unless otherwise indicated. To improve visibility of deeper tissues, embryos were treated with 0.2 mM 1-phenyl-2-thiourea (PTU) starting at 24 hpf to inhibit pigmentation. Live embryos were anesthetized with 0.016% tricaine and mounted in 0.6% low-melting-point agarose. A 20x water-immersion objective was used for most fluorescence imaging, and a 60x water-immersion objective was used for high-resolution analysis of DA geometry and endothelial cell boundaries and for counting *pdgfrb:GFP-*positive cells. The Nikon Elements AR Denoise.ai algorithm was applied to images with high background signal. Image acquisition parameters were maintained consistently within each experiment.

### Measurement of pulsatile DA diameter change

Embryos obtained from an incross of *she^ci26^* heterozygous parents in the *Tg(kdrl:GFP)* background were imaged live at 84 hpf using the resonant scanner of a Leica DMi8 confocal microscope. A single optical plane through the DA was recorded for 10 seconds using a 20x objective and 4x digital zoom. Embryos were recovered and genotyped after imaging. The minimum/contraction diameter (D_C_) and maximum/expansion diameter (D_E_) were measured using Imaris 10.0 and NIS-Elements software. Fractional diameter change was calculated as (D_E_ - D_C_)/D_C_.

### DA cross-sectional shape and size

Embryos from an incross of *she^ci26^* heterozygous parents carrying *Tg(kdrl:HRAS-mCherry)* were imaged at 3 dpf. Three-dimensional image stacks were rotated to visualize comparable DA cross-sections. Horizontal (D_H_) and vertical (D_V_) diameters were measured, and the D_H_/D_V_ ratio was calculated as an index of vessel circularity. A cross-sectional surface was generated in Imaris, and its area was measured to quantify vessel size. Two measurements of each metric were taken for each embryo, and the measurements were averaged to obtain the final estimates. All measurements were performed blinded to genotype.

### Measurement of DA diameter and glomerular capillary tuft (GCT) size

The DA diameter in MTZ-treated, SNP-treated, and *elnb* morpholino-injected embryos was measured as previously described [28]. Three measurements were taken at alternate intersegmental vessel (ISV) intervals in the trunk, with each measurement positioned midway between two adjacent ISVs. The three measurements were averaged to obtain one value per embryo for statistical analysis.

To analyze GCT size, 4 dpf embryos, obtained from an in-cross of *she^ci26^* heterozygous fish in a *Tg(fli1:lifeact-GFP)* endothelial reporter background, were fixed in 4% paraformaldehyde (PFA) for 2 hours. After washing with PBT, the yolk sac was dissected with ultrasharp tweezers to expose the GCT. We imaged the GCT using a confocal microscope and measured the area encompassing the GCT in maximum-intensity projections of each image to estimate GCT size.

### Endothelial cell shape and size

Endothelial cell geometry was analyzed in wild-type and *she^ci26^-/-* embryos carrying *Tg(kdrl:HRAS-mCherry; tjp1a:eGFP)*, which labels endothelial membranes and cell-cell junctions. The DA was imaged at 2 dpf using a 60x water-immersion objective. Cell boundaries were manually traced in three-dimensional image stacks in a region spanning two somites in the mid-trunk and computationally unrolled onto a two-dimensional surface using a previously validated method [38]. Cell elongation, orientation, surface area, and perimeter were quantified. Embryos were genotyped after imaging.

### Quantification of mural and perivascular stromal cells

To analyze mural cells and perivascular stromal cells, we crossed *she^ci26^* heterozygous fish carrying the *Tg(kdrl:HRAS-mCherry)* endothelial reporter with the *Tg(pdgfrb:GFP)* mural cell reporter to obtain double-transgenic mutant and wild-type embryos. Trunk vessels were imaged at 3 dpf. Using the spot-detection algorithm in Imaris 10.0, we separately counted *pdgfrb:GFP*-positive cells closely associated with the DA (mural cells) and cells occupying the space between the DA and posterior cardinal vein (PCV) (perivascular fibroblast-like stromal cells) within two somites in the mid-trunk. Embryos were genotyped after imaging.

### Mural cell ablation

Embryos were obtained from incrosses of *she^ci26^+/-*; *pdgfrb:gal4; UAS:mCherry-NTR; kdrl:GFP*-positive adults. After selection for mCherry and GFP expression, embryos were incubated in 50-100 µM metronidazole (MTZ, M3761, Sigma-Aldrich) or vehicle (0.1% DMSO diluted with RO grade water) control beginning at 3 dpf. Mural cell depletion was evaluated by confocal imaging at 4 dpf. At 4-5 dpf, embryos were scored for blood circulation and pericardial edema, and DA diameter was measured. Embryos were subsequently genotyped.

### Sodium nitroprusside (SNP) treatment

A fresh 1 mM SNP (LabChem, USA) stock was prepared in embryo medium and diluted to a final concentration of 1 µM. Embryos were obtained by incrossing rescued *she^ci26^*-/-; Tg*(fli1:she-2A-mCherry; kdrl:GFP)* adults. Non-rescued mCherry-negative mutant embryos and mCherry-positive rescued controls were treated with SNP beginning at 3.5 dpf and analyzed at 5 dpf. Pericardial edema and blood circulation were scored, and DA caliber was quantified by confocal imaging.

### Cardiac imaging and distensibility

Embryos from an in-cross of *she^ci26^* heterozygous parents were imaged at 4 dpf before overt pericardial edema. Brightfield movies were acquired for 10 seconds in a consistent cardiac plane using a 20x objective on a Nikon Eclipse microscope. Minimum and maximum chamber dimensions were used to calculate fractional distensibility of the atrium, ventricle, and bulbus arteriosus (BA) using the same equation applied to the DA. Diastolic two-dimensional surface areas of the atrium and ventricle were measured using NIS-Elements AR. Retrograde flow across the atrioventricular (AV) valve was evaluated from high-speed movies.

### Immunofluorescence (IF)

We used two different IF protocols. For Elnb, Collagen I pro-peptide and 3G8, 4 dpf embryos were fixed overnight at 4 °C in 4% PFA, dehydrated in 100% ethanol, rehydrated, and washed in PBT (1X PBS + 0.1% v/v Tween 20). Samples were permeabilized in 0.5% Triton X-100 in PBS for 2 hours at room temperature (RT) and blocked for 2.5 hours. Embryos were incubated overnight at 4°C with primary antibodies: rabbit anti-Elnb (1:1,000, provided by Dr. Geoffrey Burns, Boston Children’s Hospital), mouse SP1.D8 anti-Collagen I pro-peptide (1:500, Developmental Studies Hybridoma Bank [DSHB], Iowa, USA), or mouse 3G8 anti-proximal tubule (1:100, European Xenopus Resource Center, Portsmouth, UK). After washing, samples were incubated overnight with donkey anti-rabbit Alexa Fluor 647 (Elnb), donkey anti-mouse Alexa Fluor 594 (3G8), and donkey anti-mouse Alexa Fluor 647 (SP1.D8) secondary antibodies (1:500, Invitrogen). Embryos were imaged on a Nikon AX confocal microscope using a 20x water-immersion objective. Three-dimensional surfaces encompassing the Elnb signal were generated in Imaris to quantify signal volume. Integrated density of ElnB and Collagen I pro-peptide was measured from maximum-intensity projections in Fiji and normalized to the mean value of the wild-type group within each replicate.

For laminin staining, embryos were fixed in 1% PFA for 2 hours at room temperature, washed in PBT, and permeabilized in 1% Triton X-100 in PBS for 1 h at room temperature. Samples were blocked in a saponin blocking buffer (0.2% w/v Saponin + 2 mg/ml BSA + 10% v/v Lamb Serum dissolved in 1X PBS). Embryos were incubated overnight at 4°C with rabbit anti-laminin (LAMA1) antibody (1:100, L9393 Sigma-Aldrich, USA), washed, and incubated with donkey anti-rabbit Alexa Fluor 647 secondary antibody (1:500, Invitrogen) for 2 hours at room temperature. Mean laminin fluorescence intensity was measured from maximum-intensity projections in Fiji.

### *elnb* morpholino injection

A previously validated translation-blocking *elnb* morpholino [39] (5’-CCGGGCCATCCTGCTCTGTAATAAC-3’) was obtained from Gene Tools. One nanogram was injected at the one-cell stage into embryos generated by incrossing homozygous *she* mutants carrying the *Tg(fli1:she-2A-mCherry; kdrl:GFP)* rescue transgene. mCherry-positive rescued embryos and mCherry-negative *she^ci26^* mutants were separated by fluorescence screening. Elnb reduction was confirmed by immunofluorescence at 4 dpf. At 4-5 dpf, embryos were scored for circulation and pericardial edema, and DA diameter was measured.

### Whole-mount in situ hybridization and HCR

Whole-mount in situ hybridization was performed as previously described [40, 41]. DIG-labeled antisense probes were generated for *klf2a* and *notch1b.* To synthesize the *klf2a* probe, the klf2a-pExpress-1 construct (Horizon Discovery, USA) was linearized with SmaI (ThermoFisher) and transcribed using T7 RNA polymerase (Promega). To synthesize the *notch1b* probe, the *notch1b* construct was linearized with BamHI-HF (NEB) and transcribed using T3 RNA polymerase (Promega).

Fluorescent hybridization chain reaction (HCR) [42] for *col1a1, col5a1,* and *col4a1* was performed using probes (v3.0) and Alexa Fluor 594 and 647 hairpins from Molecular Instruments. Embryos from *she+/-; kdrl:GFP* incrosses were processed, imaged, and subsequently genotyped. To analyze *col4a1* expression, we generated a three-dimensional endothelial surface from the GFP channel to mask nonvascular *col4a1* signal. Mean *col4a1* fluorescence intensity and signal density surrounding a defined DA segment were quantified in Imaris. To quantify *col1a1* and *col5a1* expression, we measured the width of each expression domain above and below the DA across a three-somite region, with three measurements per somite. These measurements were averaged to determine the mean width of each expression domain. Integrated density of *col1a1* mRNA signal above the DA was also measured from maximum-intensity projections in Fiji and normalized to the mean value of the wild-type group within each replicate. All analyses were performed blinded to genotype.

### Embryo dissociation and single-cell suspension preparation for single-cell RNA sequencing

Homozygous *she^ci26^-/-* embryos were obtained from an incross of *she^ci26^-/-;fli1:she-2A-mCherry;flk1:GFP* adults, which survive to adulthood because of the rescuing *fli1:she-2A-mCherry* transgene. Wild-type siblings of these rescued adults that also carried the rescue transgene were in-crossed to generate control embryos. Embryos from both crosses were screened for mCherry expression, and only mCherry negative mutant and wild-type embryos were selected for dissociation. Approximately 30 embryos were pooled per experimental group in each biological replicate.

Cell dissociation was performed using a cold-protease dissociation protocol [43]. Embryos were transferred to 2 mL microcentrifuge tubes and triturated in 1 mL of deyolking buffer until the yolk sac was completely dissolved. Embryos were pelleted at 300x g for 1 minute, washed twice with 0.5X Danieau buffer, and resuspended in a cold protease dissociation mix containing 10 mg/mL *Bacillus licheniformis* protease (Sigma-Aldrich, USA), 125 U/mL DNase I, and 0.5 mM EDTA in 7.5% BSA (Gibco, USA). Samples were incubated on ice and triturated 15 times every 2 minutes using a 1 mL pipette tip. Dissociation was monitored microscopically every 5 minutes until a uniform single-cell suspension was achieved (approximately 20 minutes). Cells were collected by centrifugation at 1,200x g for 5 minutes at 4°C, washed, and filtered through a 20 μm cell strainer. Following two consecutive wash steps in 7.5% BSA, cell pellets were resuspended in 7.5% BSA. Cell concentration and viability were quantified using a hemocytometer.

### Single-cell RNA library preparation and sequencing

Immediately after dissociation, cells were fixed and stored using the Evercode Cell Fixation Kit (Parse Biosciences, USA) according to the manufacturer’s protocol. Approximately 21,000-22,000 fixed cells per replicate per group were barcoded and processed into whole-transcriptome single-cell RNA sequencing libraries using the Evercode Whole Transcriptome WT v3 kit (Parse Biosciences, USA) following the split-pool combinatorial barcoding workflow. Eight sublibraries were generated and sequenced by Novogene using the NovaSeq XPlus Platform (Illumina) with paired-end 150 bp reads, generating over 3.1 billion total reads at an average sequencing depth of ∼63,000 reads per cell.

### Bioinformatic processing

FASTQ files were processed using the Trailmaker platform (Parse Biosciences, USA). The integrated Parse Biosciences pipeline (v1.2.1) performed barcode demultiplexing, read alignment to the zebrafish reference genome (*Danio rerio*, GRCz11), transcript quantification, and initial cell calling using an empty-droplet classifier at a false discovery rate (FDR) threshold of 0.01 across all six biological samples, identifying an initial 49,919 cells. Library-specific cell-size filtering retained cells meeting minimum transcript thresholds ranging from 709.8 to 924.8 UMIs (unique molecular identifiers) per cell. Cells with elevated mitochondrial transcript content were excluded using an absolute threshold cutoff of ≤ 0.64%-0.72% (maxFraction = 0.0064-0.0072), and outliers with aberrant gene-to-UMI ratios were filtered via spline regression (p<0.00012-0.00016). Doublets were identified and removed using scDblFinder with sample-specific probability thresholds set between 0.820 and 0.838. After filtering, 43,144 cells were retained across all six samples (three wild-type and three mutant biological replicates), averaging 7,190 cells per sample, with medians of 2,037 detected genes and 4,853 transcripts per cell.

Downstream analyses were performed within the Trailmaker analysis suite utilizing an integrated Seurat workflow. Gene counts were normalized using standard log-normalization (logNormalize, scale factor = 10,000), and the top 2,000 highly variable genes were selected for dimensionality reduction. Reciprocal Principal Component Analysis (rPCA) was performed to calculate the top 30 principal components, followed by batch integration across samples using Harmony. Unsupervised clustering was carried out using the Leiden community detection algorithm at a resolution of 0.8, and clusters were embedded into two dimensions using UMAP with a cosine distance metric and a minimum distance of 0.3. Cluster annotation was performed using the expression of the top 50-100 marker genes with the help of ChatGPT 5.5.

To assess genotype-dependent changes in cell abundance, cluster-specific frequencies were computed in Trailmaker. Cells were grouped by genotype (wild-type vs. *she* mutant), and the relative proportion of each of the 27 Leiden clusters (resolution = 0.8) was calculated by normalizing the cell count of each cluster to the total pool of quality-filtered cells within that genotype.

To resolve vascular and stromal cell heterogeneity, highly *col1a1a*-expressing putative perivascular fibroblast, *kdrl*-expressing endothelial cell (EC), and strongly *acta2*-expressing vascular smooth muscle cell (SMC) populations were annotated and subclustered. Differences in cell abundance across genotypes were evaluated across independent biological replicates (n=3 *she+/+*, n=3 *she-/-*) using an unpaired Student’s *t*-test in GraphPad Prism, with statistical significance defined as *p* < 0.05.

Pairwise differential gene expression analysis between wild-type and mutant cells was conducted using a two-sided Wilcoxon rank-sum test with Benjamini-Hochberg false discovery rate correction; genes meeting an adjusted significance threshold of FDR ≤ 0.05 (-log10(adjusted p-value) ≥ 1.30) and an absolute effect size of log2 (fold change) ≥ 1.0 (corresponding to a minimum two-fold change in expression) were defined as significantly differentially expressed genes. UMAP plots, cell abundance frequency plots, volcano plots, and feature plots were obtained and exported from Trailmaker.

The scRNA-seq data have been submitted to NCBI GEO database and are available under accession number GSE352057.

### Statistical analysis

Statistical analyses were performed using GraphPad Prism 11. Differences between experimental groups were evaluated using Student’s t-test, one-way ANOVA, and Fisher’s exact test, as appropriate. Results are presented as mean ± standard error of the mean (SEM). All measurements were performed blinded to genotype, with embryos genotyped afterward. Each experiment was repeated at least twice with independent biological replicates.

## Results

### The enlarged DA retains normal endothelial geometry in *she* mutants

We previously showed that *she^ci26^* mutant embryos develop an enlarged DA at 1-2 dpf, followed by progressive narrowing and loss of circulation beginning at approximately 4 dpf [28] (Supplementary Movie 1). Our previous data demonstrated that increased endothelial cell number contributes to the early enlargement, but it remained unclear whether altered endothelial cell geometry also changes vessel architecture before collapse. To address this question, we analyzed endothelial cell shape in the DA at 2 dpf using the *Tg(tjp1a:eGFP)* reporter line to visualize cell-cell junctions. Endothelial cell orientation, elongation, surface area, and perimeter were not significantly different between *she* mutants and wild-type siblings (Supplementary Fig. 1).

We next examined DA cross-sectional geometry at 3 dpf, before overt circulatory failure. The cross-sectional area of the DA was increased in *she* mutants, consistent with the previously reported enlargement of the vessel (Fig. 1A-E). However, horizontal and dorsoventral diameters increased proportionally, and the D_H_/D_V_ ratio was unchanged (Fig. 1C, D, F, G). Thus, the *she* mutant DA remains approximately cylindrical, and the endothelial layer retains normal cellular organization before collapse, despite the increased vessel size.

**Figure 1.**
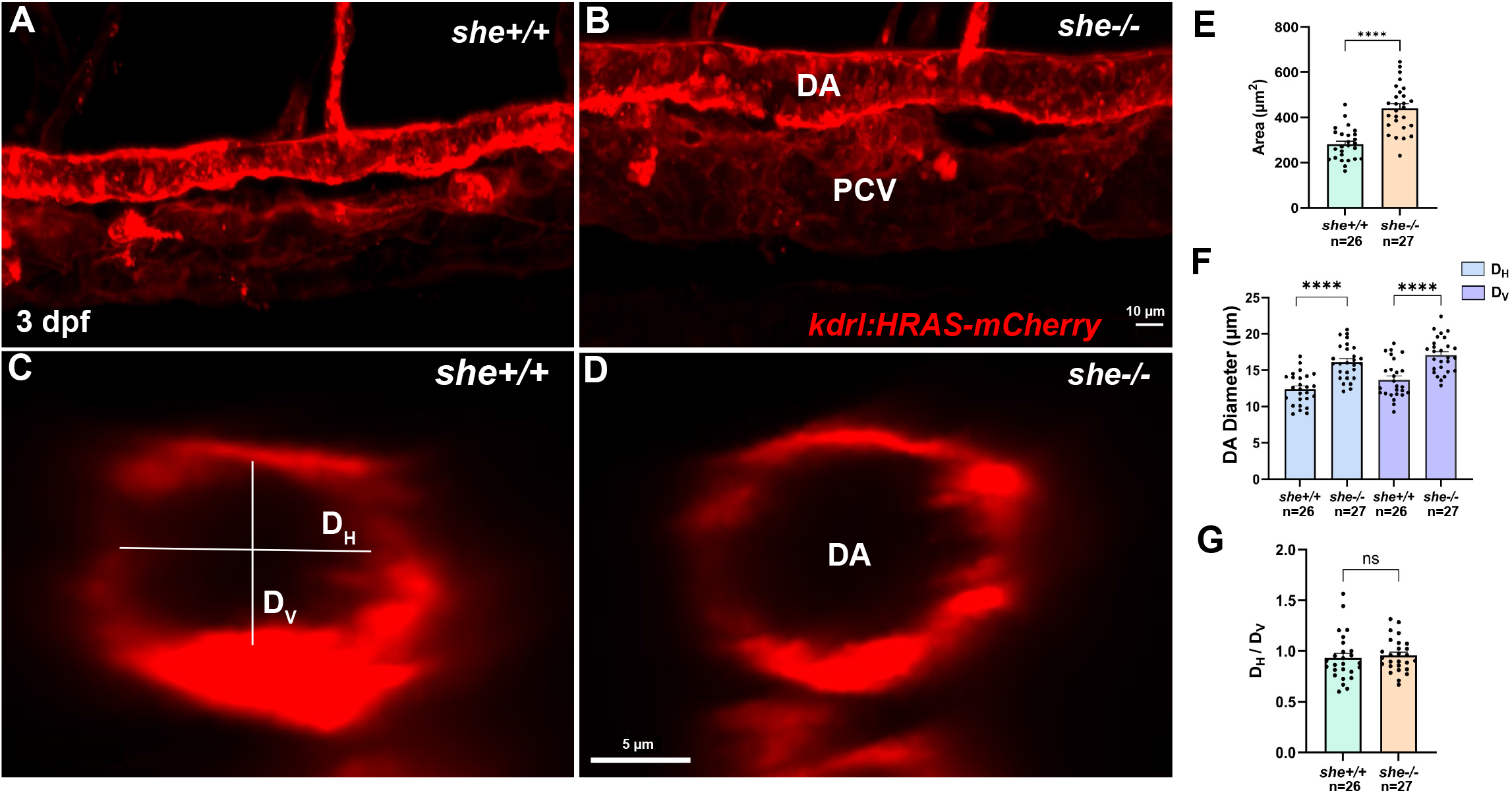
*she* mutants show increased size of the dorsal aorta (DA). (A-D) Representative confocal images of the DA in wild-type (A, C) and mutant (B, D) zebrafish embryos in the *kdrl:HRAS-mCherry* endothelial reporter background at 3 dpf in sagittal (A, B) and transverse (C, D) planes. (E) Quantification indicates a larger DA cross-sectional area in *she-/-* embryos than in *she+/+* siblings. (F) Measurement of the horizontal (D_H_) and vertical (D_V_) diameters of the DA in *she-/-* mutants and sibling wild-type embryos. (G) Quantification of DA circularity/ovality, calculated as the ratio of D_H_ to D_V_, suggests no significant change in vessel shape in *she-/-* embryos compared with *she+/+* siblings. Scale bar: 10 µm (A, B) and 5 µm (C, D); n*_she+/+_* = 26; n*_she-/-_* = 27; ns, not significant; ****p<0.0001; Student’s t-test. Error bars show SEM. Data were combined from two replicate experiments.

### She deficiency increases DA distensibility

Because DA collapse occurs after a period of apparently normal blood circulation, we asked whether arterial wall mechanical properties were altered before failure. Arterial distensibility is defined as a relative change in arterial volume or diameter per unit change in pressure and is a commonly used parameter to evaluate arterial elasticity and stiffness. Live imaging at 3.5 dpf revealed a significantly greater fractional change in DA diameter during the cardiac cycle in *she* mutants than in wild-type siblings (Fig. 2; Supplementary Movie 2). These data suggest that the mutant DA is more distensible and therefore less resistant to pulsatile deformation.

**Figure 2.**
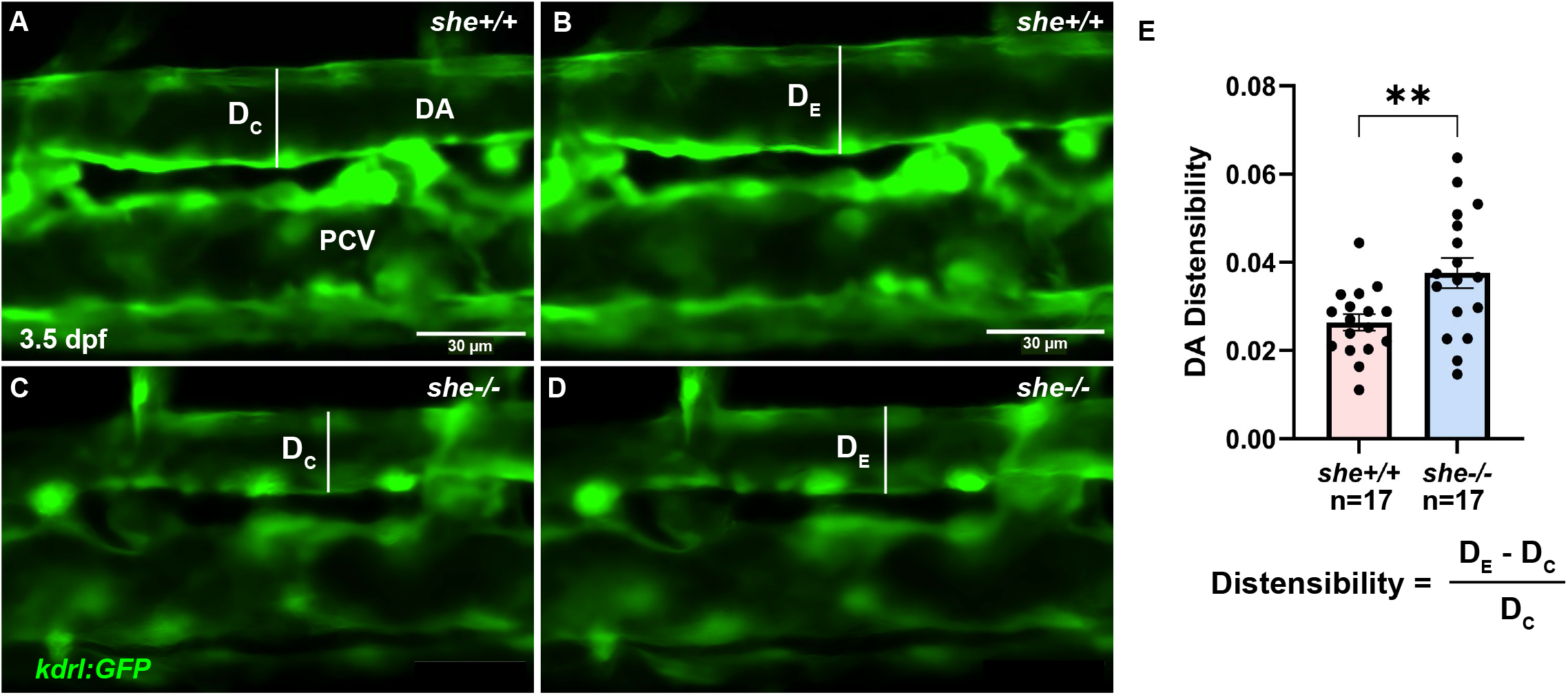
Loss of She leads to increased distensibility of the DA in developing zebrafish embryos. (A-D) Representative time-lapse stills from confocal imaging of wild-type (A, B) and *she-/-* mutant (C, D) live zebrafish embryos expressing the *kdrl:GFP* endothelial reporter, capturing the movement of the DA wall as it expands (B, D) and contracts (A, C) with each heartbeat. (E) Quantification of DA distensibility, calculated as the fractional change in vessel diameter [(Expansion diameter {D_E_} – Contraction diameter {D_C_})/D_C_]. Scale bar: 30 µm; n*_she+/+_* = 17; n*_she-/-_* = 17; **p<0.01; Student’s t-test. Data were combined from two independent experiments.

### *she* mutants show reduced abundance of perivascular stromal cells

Mural cells progressively associate with the zebrafish DA between 2 and 5 dpf, overlapping temporally with the onset of DA narrowing in *she* mutants [14, 25]. We therefore examined whether mural cell abundance was altered in *she* mutants. Quantification of *Tg(pdgfrb:GFP)*-positive cells associated with the DA at 3 dpf revealed no significant difference between *she* mutants and wild-type siblings (Fig. 3A-C), suggesting that the changes in DA distensibility or subsequent DA collapse are not accompanied by a detectable change in DA-associated mural cell number.

**Figure 3.**
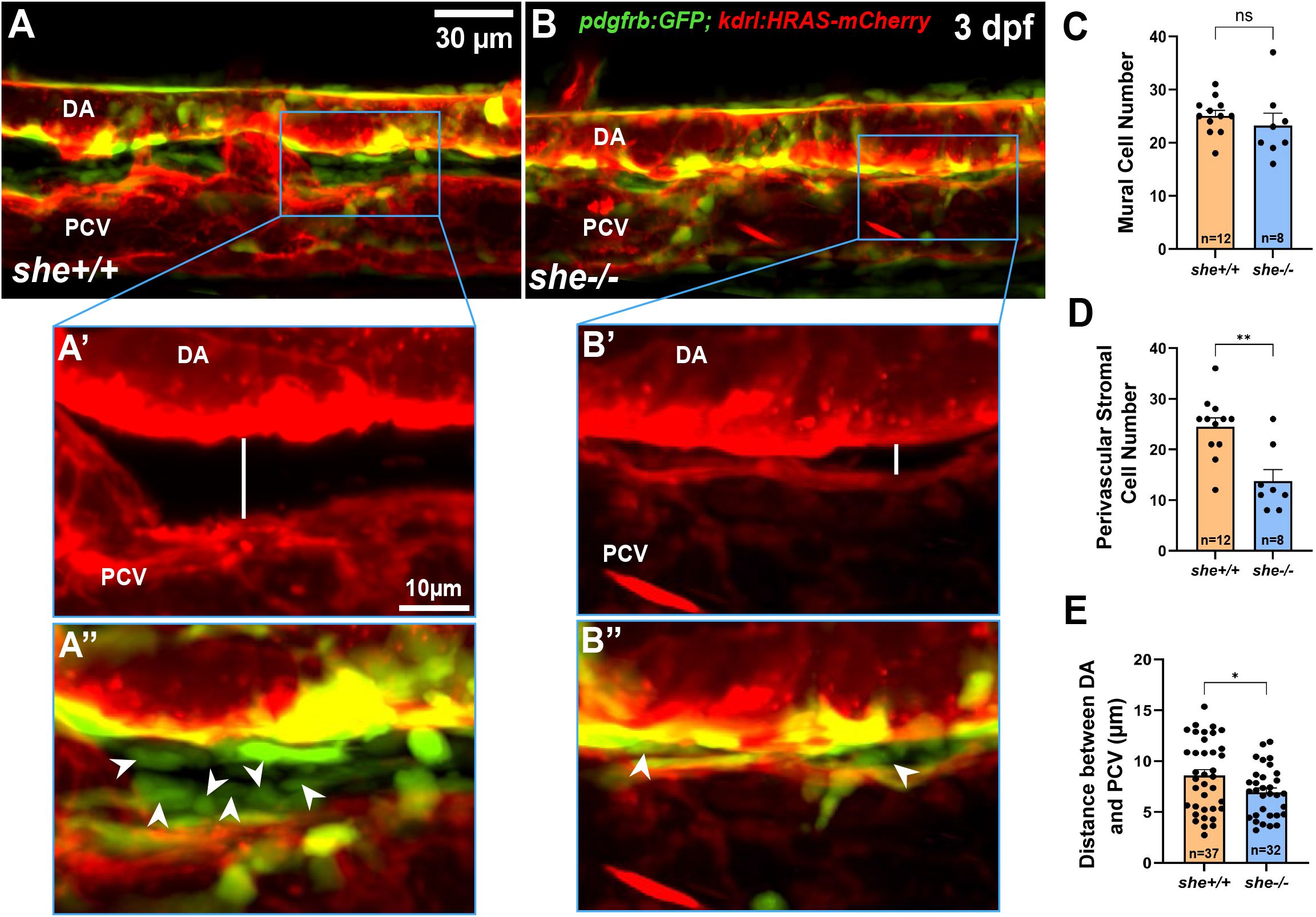
The number of perivascular stromal cells is reduced in *she* mutants. (A-B”) Representative confocal images of the trunk vasculature in wild-type (A-A”) and *she-/-* mutant (B-B”) zebrafish embryos in the *Tg(pdgfrb:GFP; kdrl:HRAS-mCherry)* reporter background at 3 dpf. *pdgfrb:GFP* (green) labels the mural cells surrounding the DA, while *kdrl:HRAS-mCherry* (red) labels the endothelial cell membranes. (A, B) Merged confocal projections showing an overview of Pdgfrb-positive cells associated with the trunk vasculature; boxed regions are magnified in (A’-B”). (A’-B’) Single-channel (*kdrl:HRAS-mCherry*) magnified images showing the intervessel space between the DA and PCV in *she+/+* (A’) and *she-/-* (B’) embryos. Note the reduced distance between the DA and PCV in *she* mutant embryos. (A”-B”) Merged magnified images showing the differential abundance of low *pdgfrb:GFP*-expressing cells in the intervessel space of *she+/+* (A”) and *she-/-* (B”) embryos. Note the reduced number of *pdgfrb:GFP* cells in *she* mutants. (C-E) Quantification of high *pdgfrb:GFP*-expressing mural cells associated with the DA (C), the low *pdgfrb:GFP*-expressing perivascular stromal cells (D), and the distance between the DA and PCV (E). Horizontal white lines indicate the intervessel space between the DA and PCV. Scale bar: 30 µm (A, B) and 15 µm (A’-B’’). n*_she+/+_* = 12 (C, D) and 37 (E); n*_she-/-_* = 8 (C, D) and 32 (E); ns, not significant; *p<0.05; **p<0.01; Student’s t-test. Data were combined from three independent experiments.

The DA and PCV are located in close proximity at 1-2 dpf stage during normal development of wild-type embryos. However, the distance between the DA and PCV substantially increases between 3 and 5 dpf [44, 45]. Intriguingly, the gap between the DA and PCV was significantly narrower in *she* mutant embryos compared to wild-type siblings at 3 dpf (Fig. 3A’, B’, E). The *Tg*(*pdgfrb*:GFP) reporter labeled a population of weakly fluorescent stromal cells in this space (Fig. 3A”, B”), and their number was significantly reduced in *she* mutant embryos compared to wild-type siblings (Fig. 3D).

Unlike DA-associated mural cells, these weakly *pdgfrb*:GFP-positive cells did not closely adhere to the axial vessels. *Pdgfrb* expression is known to label several different cell populations, including perivascular stromal cells, which express multiple different collagen genes [46, 47]. To verify their identity, we conducted HCR for *col1a1* in wild-type *Tg(pdgfrb:GFP)* embryos. Most of these low *pdgfrb:GFP*-positive cells expressed *col1a1* (Supplementary Figure 2D-I), supporting their identification as perivascular fibroblasts. Some *col1a1*-positive cells lacked detectable GFP, indicating heterogeneity within this fibroblast population (Supplementary Figure 2D-F). *col1a1* expression was also observed in DA-associated mural cells and in perivascular cells above the DA.

Stromal cells contribute to vascular ECM production, including collagen and elastin [47]. This matrix normally acts as a mechanical buffer, absorbing and safely distributing the active external contractile forces generated by mural cells [18, 48]. We tested whether reduced perivascular stromal cell abundance in the mutant embryos was accompanied by altered fibrillar collagen-gene expression. HCR analysis showed near-perfect colocalization of *col1a1* and *col5a1* in perivascular cells around the DA (Figure 4A-B”). At 4 dpf, their expression domains above and below the DA were significantly smaller in mutants than in wild-type sibling embryos (Figure 4A-B”, E, F). Additionally, the integrated density of *col1a1* transcript in the domain above the DA was significantly reduced in *she* mutant embryos compared to wild-type siblings (Figure 4G). A similar reduction in the integrated density of Collagen I pro-peptide in *she* mutants was observed using whole-mount immunofluorescence (Figure 4C,D,H). Neither *elna* nor *elnb* transcripts were detected in the trunk under the conditions used. Together, these results suggest that She regulates the number of perivascular fibroblast like stromal cells surrounding the DA and in between the DA and PCV, thereby affecting fibrillar collagen expression, which likely results in reduced perivascular matrix support. Inadequate perivascular support may render the developing DA susceptible to contraction by mural cells.

**Figure 4.**
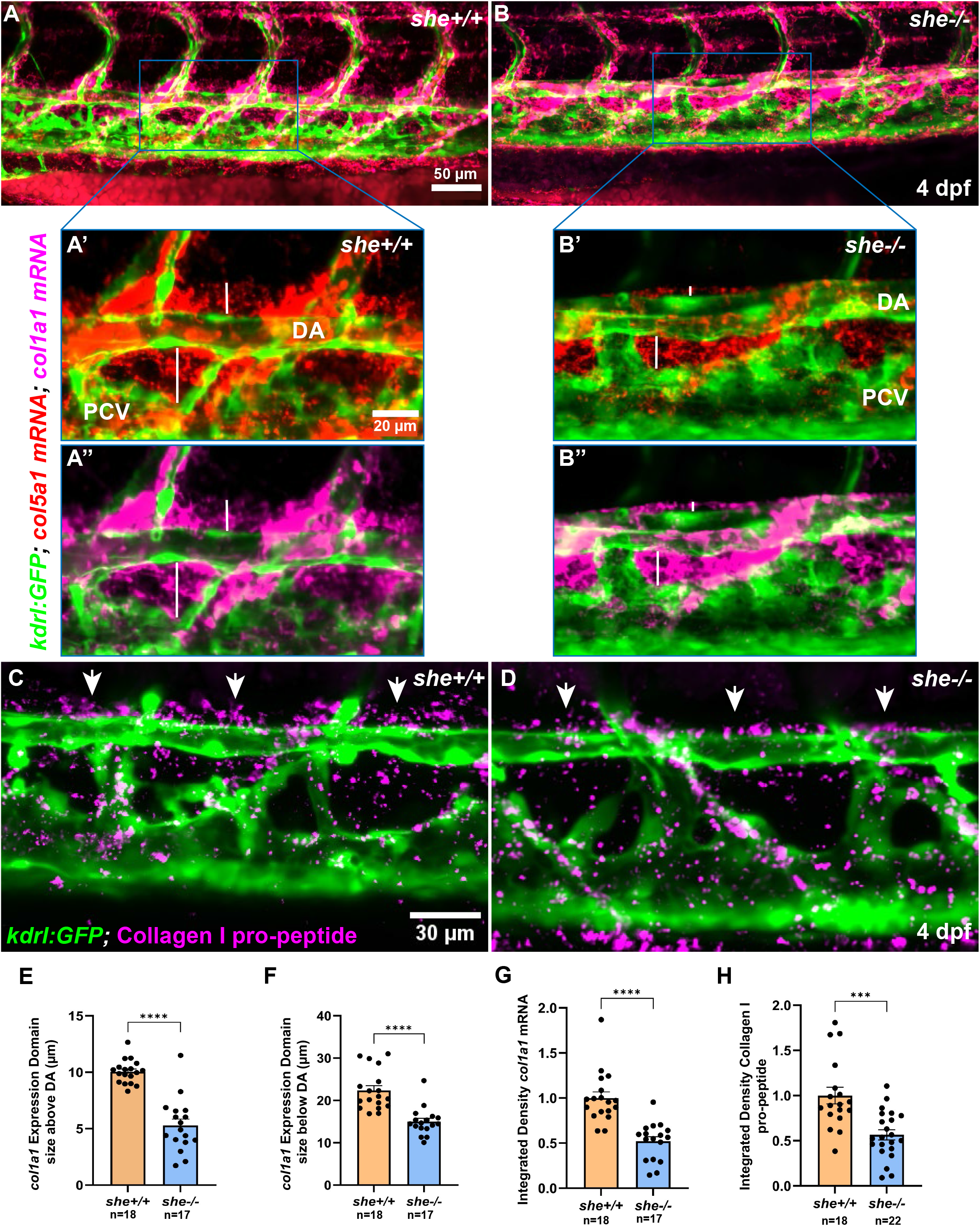
*she* mutants show reduced Collagen I expression. (A-B”) Representative confocal images of the trunk vasculature (*kdrl:GFP*; green) of wild-type (A-A”) and *she-/-* mutant (B-B”) embryos stained for *col1a1* (magenta) and *col5a1* (red) using HCR at 4 dpf. (A, B) Merged overview images of the trunk region of *she+/+* (A) and *she-/-* (B) embryos at 4 dpf. (A’, B’) Magnified boxed areas in (A) and (B), respectively, showing merged *col5a1* HCR (red) and *kdrl:GFP* (green) channels. (A’’, B’’) Magnified boxed areas in (A) and (B), respectively, showing merged *col1a1* HCR (magenta) and *kdrl:GFP* (green) channels. The vertical white lines indicate the width of *col5a1* (A’, B’) and *col1a1* (A’’, B’’) expression domains above and below the DA in *she+/+* (A’, A’’) and *she-/-* (B’, B’’) embryos at 4 dpf. Note that *col1a1* and *col5a1* expression domains largely overlap and are narrower in *she* mutant embryos than in siblings. (C, D) Representative confocal projections of the trunk vessels (green) in 4 dpf *she+/+* (C) and *she-/-* (D) embryos immunostained with SP1.D8 antibody to detect Collagen I pro-peptide (magenta). White arrows point toward Collagen I pro-peptide immunostaining in three adjacent somites. Vertical white lines indicate differences in expression-domain size above DA between *she+/+* and *she-/-* embryos. (E, F) Quantification of the size of *col1a1* expression domains above (E) and below (F) the DA in *she*+/+ (n=18) and *she-/-* (n=17) embryos. (G) Integrated density of the *col1a1* transcript signal above the DA in *she*+/+ (n=18) and *she-/-* (n=17) embryos. (H) Integrated density of immunostained Collagen I pro-peptide above the DA in *she*+/+ (n=18) and *she-/-* (n=22) embryos. Scale bar: 50 µm (A, B), 20 µm (A’-B’’) and 30 µm (C,D). ****p<0.0001, ***p<0.001; Student’s t-test. Data were combined from two replicate experiments.

### The basement membrane remains unaffected in *she* mutant embryos

Defects in the basement membrane can cause or significantly contribute to DA hyperdistensibility and subsequent vessel collapse [49–51]. While the interstitial collagen produced by fibroblasts provides bulk tissue resistance [18, 47], the basement membrane (BM), composed primarily of type IV Collagen, laminins, nidogen, and perlecan, also plays a critical role in the structural stability and elasticity of the DA [17, 52]. To assay the formation of the BM, we performed laminin IF in 4 dpf zebrafish embryos in the *kdrl:GFP* vascular reporter background. Laminin deposition was not significantly affected in *she* mutant embryos compared to wild-type siblings (Supplementary Figure 3). Similarly, *col4a1* transcript levels in the vascular region adjacent to the DA were not significantly affected (Supplementary Figure 4). These results suggest that the basement membrane is not affected in *she* mutants.

### Single-cell RNA sequencing data indicated a reduced abundance of perivascular stromal cells in *she* mutants

To assess cellular and transcriptional changes associated with She deficiency, we performed single-cell RNA sequencing on 4 dpf wild-type and *she* mutant embryos. UMAP embedding revealed 27 distinct cell clusters across both wild-type and mutant embryos (Fig. 5A). These cell clusters were annotated based on the expression of top marker genes (Supplementary Table 1). None of these clusters differed significantly in abundance between mutants and wild-type embryos (Fig. 5B). Analysis across all clusters identified 12 differentially expressed genes, with 6 upregulated and 6 downregulated in mutants. However, none of these genes showed any apparent relevance to cardiovascular development and function (Fig. 5C). In the UMAP, we annotated a smooth muscle cell population by robust *acta2* expression and a putative fibroblast population by high *col1a1a* expression (Fig. 5D, F). By subclustering, we identified a discrete fibroblast subpopulation exhibiting prominent expression of *col1a1a, col1a1b, col1a2, col5a1, fgfr2,* and other marker genes which are characteristic of fibroblasts (Fig. 5E, G, H; Supplementary Table 2). The number of these stromal cells was markedly reduced in *she* mutant samples compared with wild-type samples (Fig. 5J). Notably, the abundance of ECs and SMCs was not affected in the mutants compared to wild-type embryos (Fig. 5I, K). These findings are consistent with the reduction in perivascular stromal cells observed by imaging.

**Figure 5.**
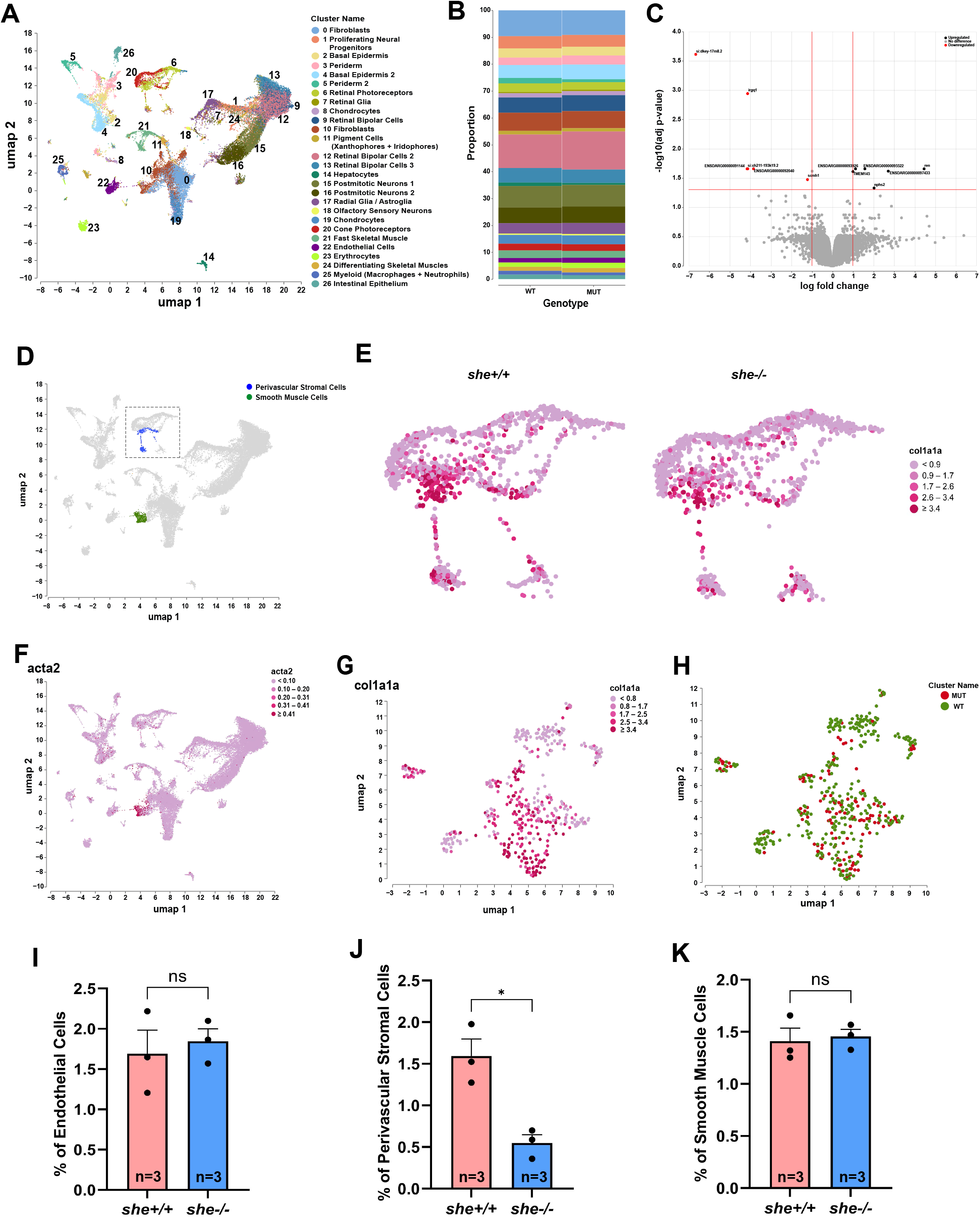
Single-cell RNA sequencing suggests a reduction in PVF cell abundance in *she* mutant embryos. (A) UMAP visualization of embryonic cell clusters resolved into 27 major cell types. (B) Proportional stacked bar plot showing relative cell-type abundance in wild-type and mutant samples. (C) Volcano plot depicting differential gene expression between wild-type and *she* mutant cells. (D) UMAP embedding of vascular mural and stromal cell subsets, with the dashed box highlighting perivascular stromal cells (blue) and smooth muscle cells (SMCs; green). (E) Feature plots showing *col1a1a-*expressing cells in wild-type (*she+/+*, left) and mutant (*she-/-*, right) samples within the region of stromal cells marked by the dashed box in (D). (F) Feature plot showing normalized expression of the smooth muscle cell marker *acta2* across all clusters. (G) Feature plot showing *col1a1a* expression levels across the subclustered perivascular stromal cell population. (H) Subclustered UMAP projection of the isolated perivascular stromal cell population, color-coded by embryo genotype (MUT, red; WT, green). (I–K) Quantification of endothelial, SMC, and PVF cell population frequencies across independent biological replicates. n*_she+/+_* = 3; n*_she-/-_* = 3; ns, not significant; *p<0.05; Student’s t-test.

### Mural cell ablation prevents DA collapse and circulation failure

Previous studies have shown that mural cells associate with the DA starting at 2 dpf and greatly increase in number at 3-5 dpf [14]. This overlaps with the time period when the DA narrows and collapses in *she* mutants at 4 dpf. We hypothesized that the DA collapse and failure of blood circulation in *she* mutant embryos could be caused by the mural cells, which initiate constriction and exert force on the wall of the DA. To test this hypothesis, we performed ablation of mural cells using the nitroreductase (NTR)–metronidazole (MTZ) approach. *she* mutants were crossed into the *pdgfrb:gal4; UAS:mCherry-NTR* transgenic line, which expresses nitroreductase (NTR) under the control of the mural cell-specific *pdgfrb:gal4* transgene. Addition of metronidazole (MTZ) at 3 dpf resulted in a reduction of mural cell coverage at 4 dpf (Fig. 6A-D). Approximately 23% of control DMSO-treated embryos, obtained from the incross of *she+/-; pdgfrb:gal4; UAS:mCherry-NTR, kdrl:GFP-*positive parents, developed pericardial edema and loss of circulation by 4-5 dpf, consistent with the expected frequency of homozygous mutants. In contrast, fewer than 4% of MTZ-treated embryos developed these defects (Fig. 6E). Genotyping of MTZ-treated embryos with preserved circulation identified 9 *she-/-* mutants among 35 embryos (26%), whereas no *she-/-* mutants were recovered among 23 vehicle-treated embryos with normal circulation. DMSO-treated *she* mutants showed the expected reduction in DA diameter at 4 dpf, whereas MTZ-treated *she* mutants retained an enlarged DA (Fig. 6F). These findings suggest that *pdgfrb-*positive mural cells contribute to the DA collapse and circulatory failure in *she* mutants.

**Figure 6.**
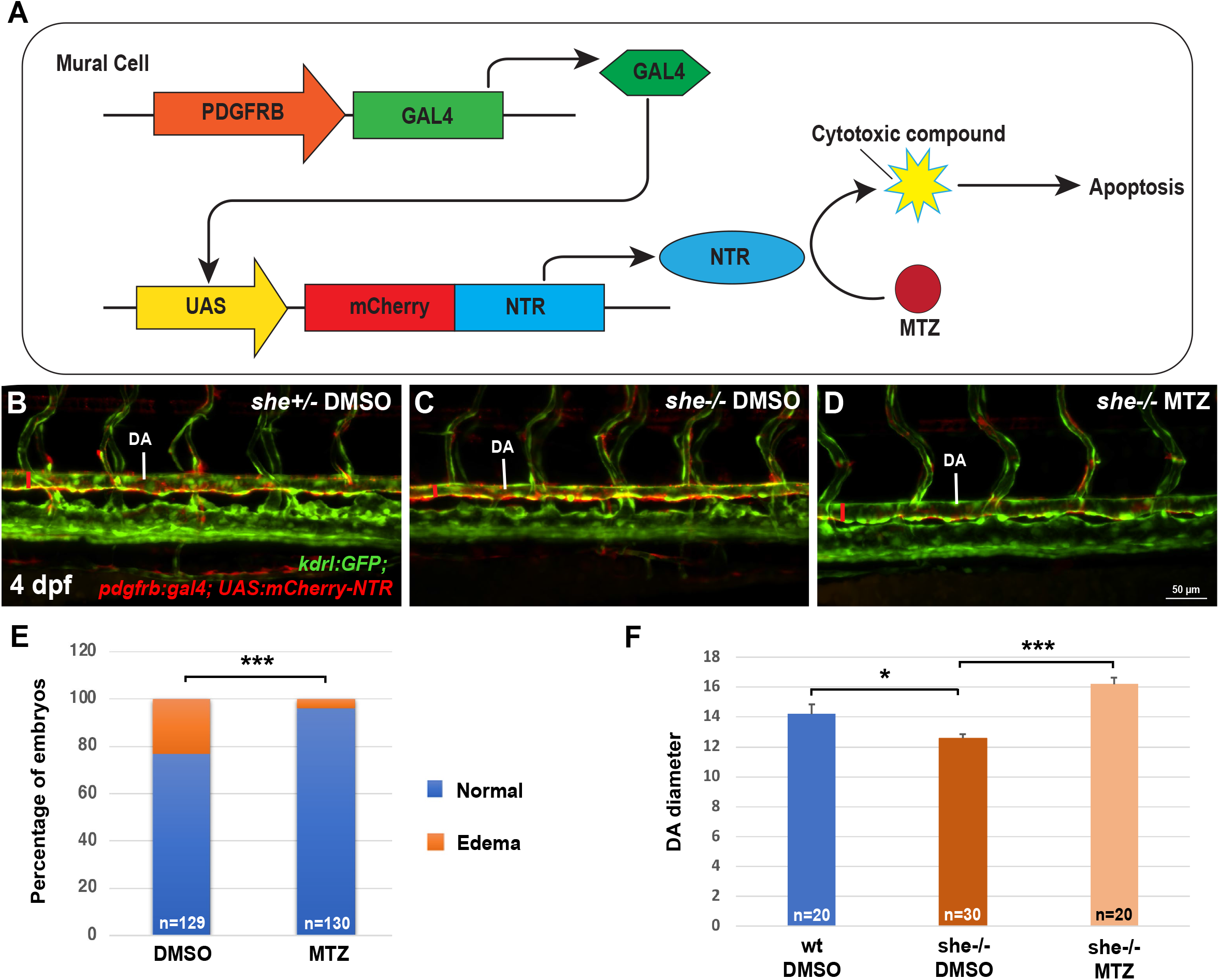
Ablation of mural cells prevents DA collapse and circulation failure in *she* mutants. (A) Schematic illustrating the UAS/Gal4-based genetic technique used to ablate mural cells. Embryos were obtained from *she+/-; pdgfrb:gal4; UAS:mCherry-NTR; kdrl:GFP* parents, sorted for mCherry and GFP expression, and treated with either MTZ or 0.1% DMSO solution at 3 dpf. NTR is expressed in *pdgfrb+* mural cells and converts MTZ into a potent toxin, leading to cell death. (B-D) Confocal images showing merged *kdrl:GFP* (green) and *pdgfrb:gal4;UAS:mCherry-NTR* (red) channels in DMSO-treated *she+/-* (B), DMSO-treated *she-/-* (C), and MTZ-treated *she-/-* (D) embryos at 4 dpf. Note the reduced mural cell coverage and wider DA in MTZ-treated *she-/-* embryos (D) compared to DMSO-treated *she-/-* embryos (C). (E) Quantification of embryos with edema and failed blood circulation at 4-5 dpf in the DMSO (23%; n=129) and MTZ treated groups (4%; n=130). Note that 25 % of embryos from this cross are expected to be homozygous *she-/-* mutants. Scale bar: 50 µm. *** p<0.001, Fisher’s exact test. (F) Quantification of DA diameter at 4 dpf in DMSO treated wild-type (n=20) and *she* mutant embryos treated with MTZ (n=20) or DMSO (n=30). Embryos were genotyped after imaging. *p<0.05, ***p<0.001, Student’s t-test.

### Pharmacological vasodilation partially rescues DA caliber and circulation in she mutants

The mural cell ablation experiment established a functional contribution of mural cells but did not distinguish their contractile activity from other mural cell functions. We therefore treated *she* mutants with SNP, an NO donor that promotes vascular smooth muscle relaxation through cGMP signaling [53, 54]. Embryos for this experiment were collected from a previously generated rescued *she* mutant line (*she-/-; fli1:she-2A-mCherry*), in which endothelial-specific expression of She completely rescues the mutant phenotype, enabling them to survive to adulthood [28]. A low SNP concentration that did not significantly alter the DA in rescued mCherry-positive controls increased DA caliber in mCherry-negative *she* mutants (Fig. 7A-E). SNP treatment also reduced the frequency of circulation failure from approximately 79% in untreated mutants to less than 60% in treated mutants (Fig. 7F). The partial rescue produced by SNP is consistent with a contribution of mural cell-dependent vascular tone to DA collapse, although systemic effects of NO signaling cannot be excluded.

**Figure 7.**
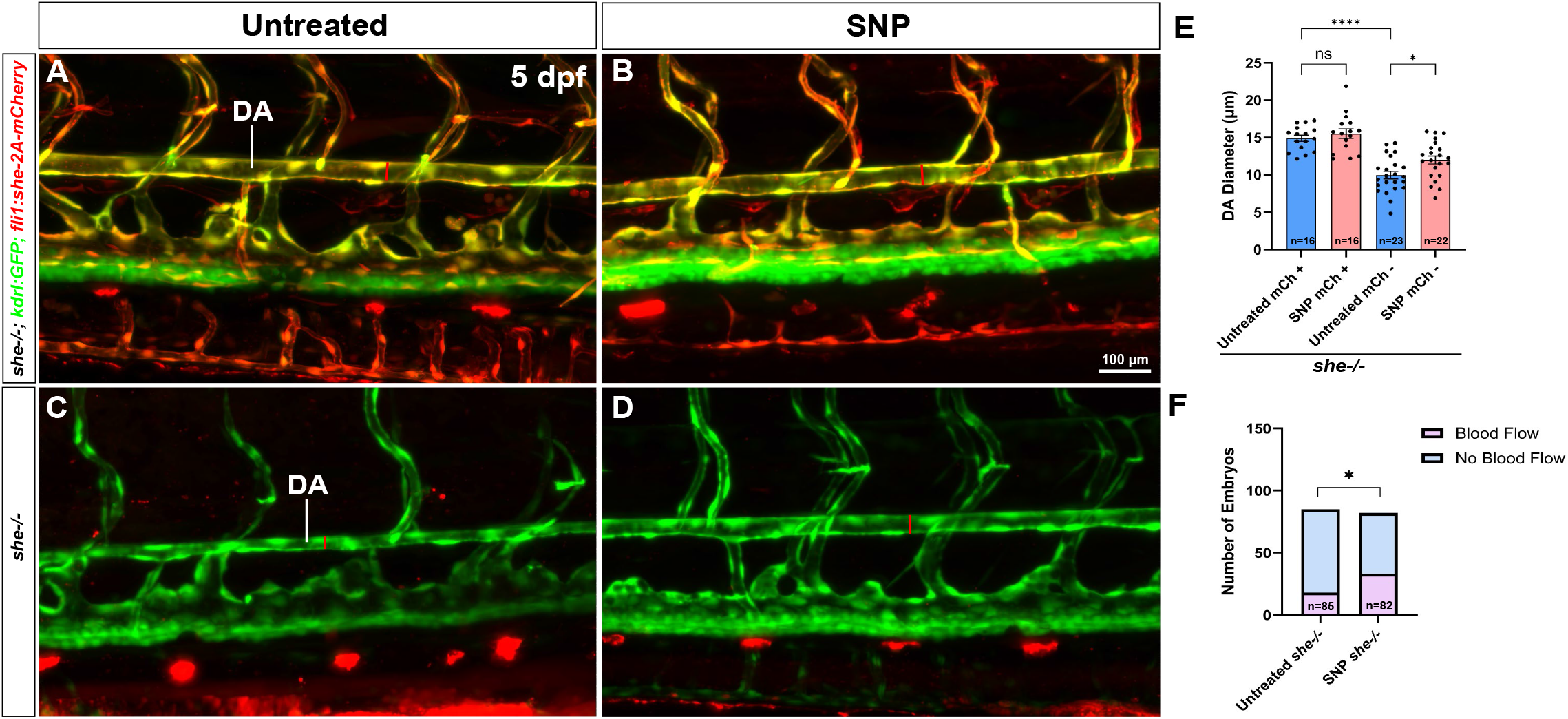
SNP-induced vasodilation partially rescues vessel collapse and blood circulation failure in *she* mutant embryos. (A-D) Confocal images showing DA diameter changes in untreated (A, C) and SNP-treated (B, D) *she-/-; fli1:she-2A-mCherry* embryos (phenotypically normal; A, B) and their sibling mCherry-negative *she-/-* embryos (C, D) in the *kdrl:GFP* reporter background (green). (E) Quantification of DA diameter. In mCherry-positive control embryos, the low SNP dose does not significantly dilate the DA, whereas the same dose rescues DA size in mCherry-negative *she-/-* mutant embryos. Data were combined from two independent experiments. n_Untreated mCh+_=16, n_SNP mCh+_=16, n_Untreated mCh-_=23, n_SNP mCh-_=22; ns, not significant, *p<0.05, ****p<0.0001, one-way ANOVA followed by Šidák’s multiple comparisons test. (F) Quantification of embryos with edema and circulatory failure at 5 dpf in untreated and SNP-treated mCherry-negative *she* mutant embryos. Scale bar: 100 µm. n_Untreated_ *_she-/-_*=85, n_SNP_ *_she-/-_*=82; *p<0.05; Fisher’s exact test.

### *she* mutants exhibit cardiac chamber dilation, altered distensibility, and atrioventricular regurgitation

Because circulatory failure may also be associated with cardiac dysfunction, we examined heart structure and function at 4 dpf. *she* mutants exhibited significantly increased diastolic areas of both the atrium and ventricle (Fig. 8L, M). High-speed imaging also showed increased fractional distensibility of the atrium and bulbus arteriosus, whereas ventricular distensibility was not significantly changed (Fig. 8A-K; Supplementary Movie 3).

**Figure 8.**
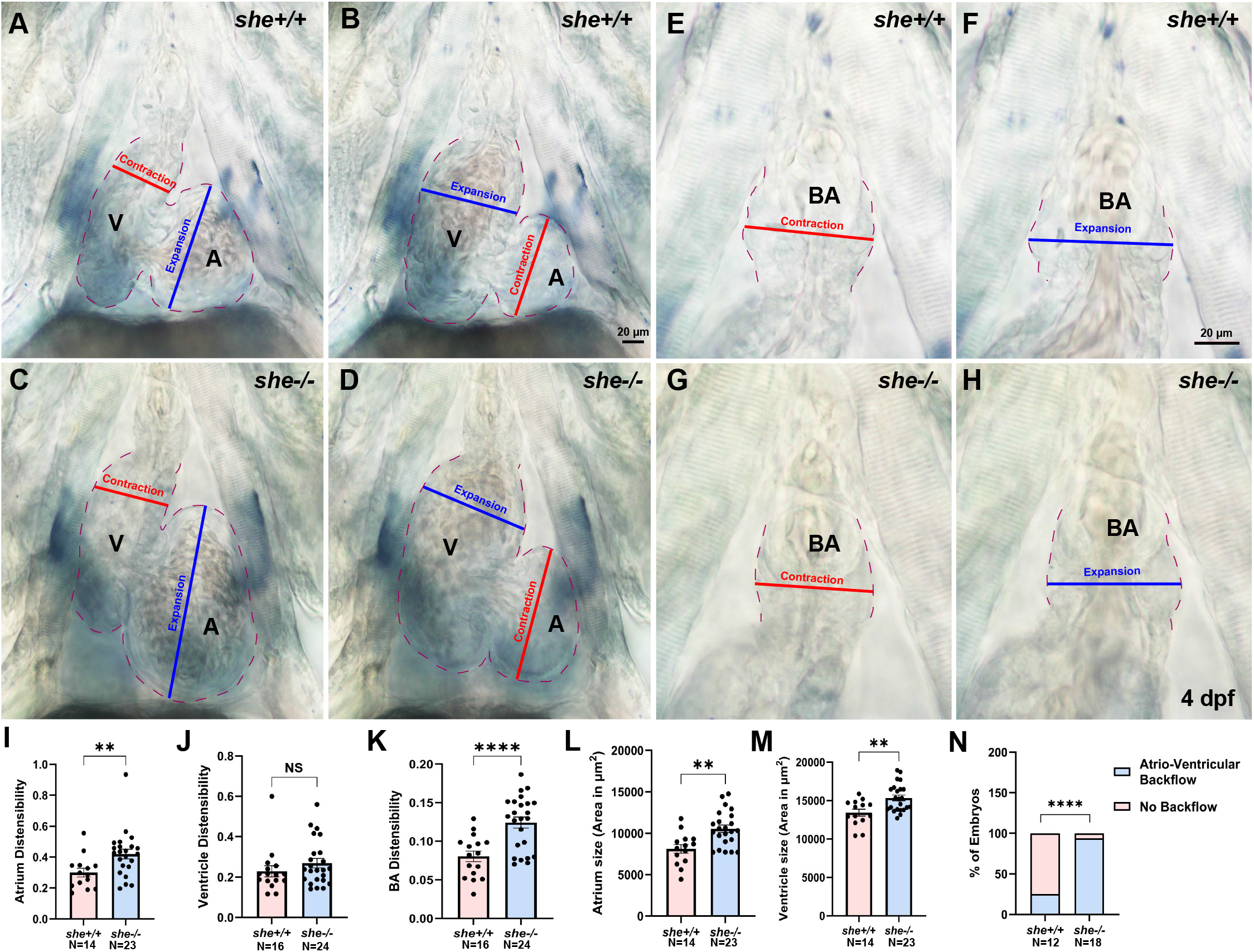
*she* mutants display defects in heart morphology, mechanics, and function. (A-H) Representative still images from Supplementary movie 3 showing the contracted and expanded heart chambers (atrium and ventricle; A-D) and outflow tract (E-H) in a single z-plane. (I-K) Quantification of distensibility in the atrium (I; n*_she+/+_* = 14; n*_she-/-_* = 23), ventricle (J; n*_she+/+_* = 16; n*_she-/-_* = 24), and BA (K; n*_she+/+_* = 16; n*_she-/-_* = 24). Before the onset of pericardial edema at 4 dpf, the atrium and bulbus arteriosus (BA) exhibit significantly greater distensibility in *she* mutants than in wild-type siblings, whereas ventricular distensibility remains unchanged. Scale bar: 20 µm. NS, not significant, **p<0.01, ****p<0.0001, Student’s t-test. (L, M) Measurements of the two-dimensional area of the atrium (L; n*_she+/+_* = 14; n*_she-/-_* = 23) and ventricle (M; n*_she+/+_* = 14; n*_she-/-_* = 23) at diastole show that both the atrium and ventricle are significantly dilated in mutants. **p<0.01, Student’s t-test. (N) At the onset of pericardial edema and circulatory failure, beginning at 4 dpf, time-lapse imaging (Suppl. movie 4) shows increased retrograde flow from the ventricle to the atrium across the AV valve in mutants compared to wild-type siblings. n*_she+/+_* =12; n*_she-/-_* =18; ****p<0.0001, Fisher’s exact test. All data shown in this figure were combined from two independent experiments.

Mutants also displayed increased retrograde blood flow from the ventricle to the atrium across the atrioventricular valve (Fig. 8N; Supplementary Movie 4). Expression of the valve-associated markers *klf2a* and *notch1b* was comparable between *she* mutants and wild-type siblings (Supplementary Fig. 5), arguing against a major defect in early valve specification. The regurgitation could arise from chamber dilation, altered loading, or abnormal valve mechanics. Because cardiac imaging was performed near the onset of DA collapse, these data do not establish whether the cardiac abnormalities precede vascular failure or arise in response to altered systemic hemodynamics.

### Elnb deposition and bulbus arteriosus size are increased in *she* mutants but do not drive DA collapse

Because the DA in *she* mutants exhibited increased distensibility, we tested whether this could be associated with altered deposition of elastin. Although immunostaining for Elnb failed to detect any apparent staining in the DA at 4 dpf, increased Elnb staining in the bulbus arteriosus was observed in *she* mutants (Fig. 9A-F). The bulbus arteriosus functions as an elastic reservoir that dampens pulsatile ventricular output [55, 56]. Endothelial and endocardial expression of She from the *Tg(fli1:she-2A-mCherry)* transgene restored Elnb signal toward control levels in *she* mutants but did not further reduce Elnb in wild-type embryos (Fig. 9G-L). These results indicate that She is required to limit excessive Elnb accumulation in the mutant background.

**Figure 9.**
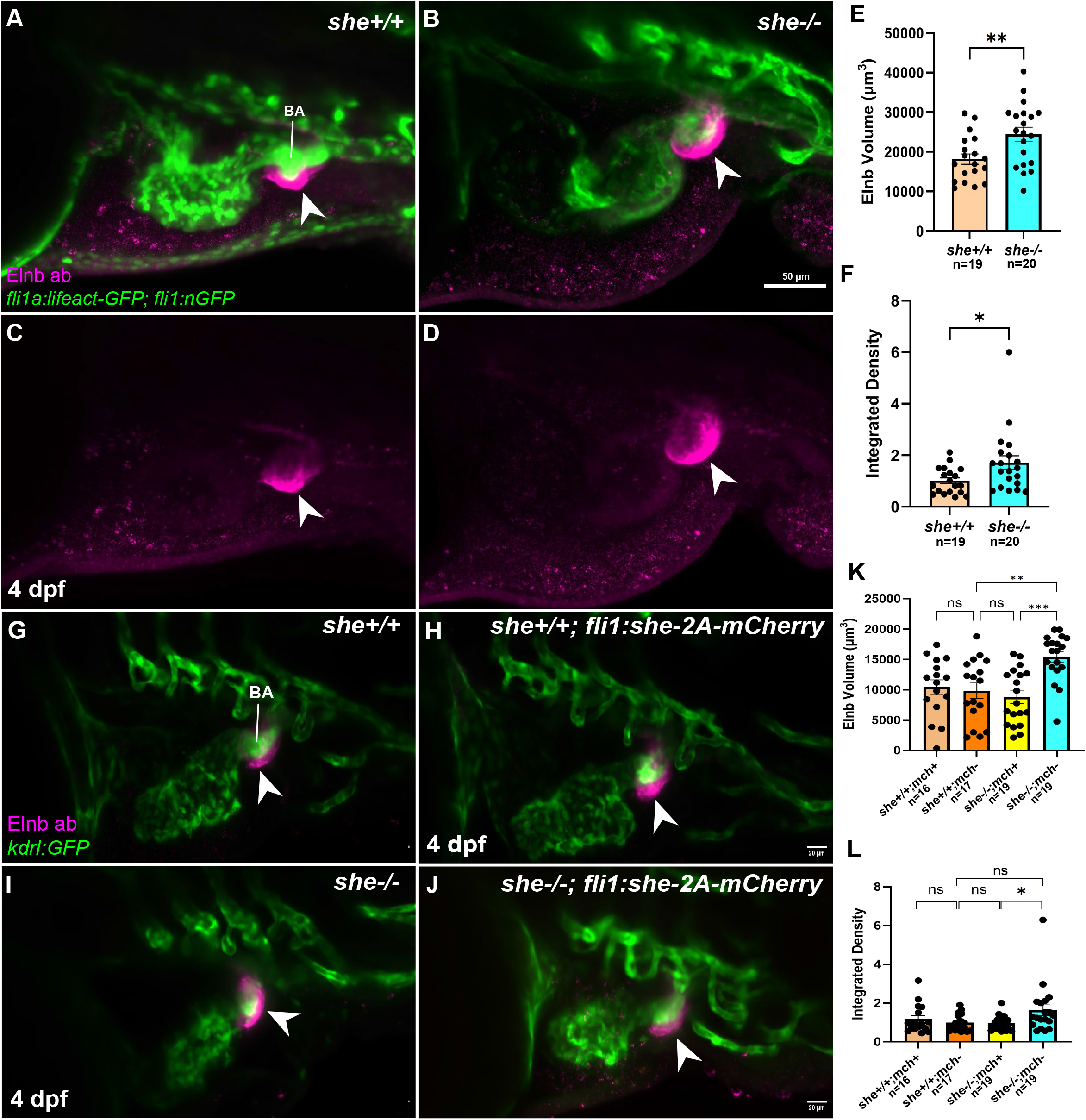
She modulates Elnb deposition around the outflow tract. (A-D) Confocal images of whole-mount immunostaining for Elnb (arrowheads; magenta) in *she+/+* (A, C) and *she-/-* (B, D) embryos obtained from an incross of *she+/-* heterozygous parents carrying endothelial markers *fli1a:lifeact-GFP* and/or *fli1:nGFP* (green) at 4 dpf. Elnb expression is localized prominently around the outflow tract and is significantly elevated in *she-/-* mutants compared to *she+/+* embryos. (E, F) Quantification of the 3D volume of Elnb signal using Imaris (E) and the normalized integrated density of the Elnb signal using Fiji (F). Scale bar: 50 µm. n*_she+/+_* =19; n*_she-/-_* =20; *p<0.05, **p<0.01, Student’s t-test. Data were combined from three biological replicates. (G-J) Confocal images showing Elnb whole-mount immunostaining (magenta) in wild-type (G, H) and mutant (I, J) embryos expressing (H, J) or not expressing (G, I) the rescue transgene *fli1:she-2A-mCherry* in the *kdrl:GFP* endothelial reporter background (green) at 4 dpf. Elnb staining is marked with an arrowhead in each image. (K, L) Quantification of the 3D volume of Elnb signal using Imaris (K) and the normalized integrated density of the Elnb signal using Fiji (L). Expression of the *fli1:she-2A-mCherry* transgene did not alter Elnb signal volume or integrated density in wild-type embryos but significantly reduced them in *she-/-* mutants compared to mCherry-negative *she-/-* mutant controls (K, L). Scale bar: 20 µm. n*_she+/+;mch+_*=16, n*_she+/+;mch-_*=17, n*_she-/-;mch+_*=19, n*_she-/-;mch-_*=19; ns: not significant, *p<0.05, **p<0.01, ***p<0.001, one-way ANOVA followed by Šidák’s multiple comparisons test. Data were combined from two biological replicates.

Consistent with increased outflow tract distensibility, three-dimensional volumetric analysis revealed enlargement of the bulbus arteriosus in *she* mutants (Supplementary Fig. 6A-C). The *fli1:she-2A-mCherry* transgene rescued this enlargement in *she* mutants but did not alter bulbus arteriosus size in wild-type embryos (Supplementary Fig. 6D-H).

To test whether increased Elnb contributes causally to systemic circulatory failure, we reduced Elnb using a previously validated translation-blocking morpholino [39]. Although Elnb immunoreactivity was reduced, Elnb knockdown did not significantly rescue DA diameter and produced only a modest, nonsignificant reduction in pericardial edema and circulation failure in *she* mutants (Supplementary Fig. 7). Thus, reducing Elnb under these conditions was insufficient to restore vascular function. Increased Elnb deposition and bulbus arteriosus enlargement are therefore unlikely to be the primary cause of DA collapse. They may represent a parallel consequence of She deficiency or a compensatory response to abnormal mechanical loading.

### Endocardial *she* expression is insufficient to rescue the mutant phenotype

Because *she* is expressed in both vascular endothelium and the endocardium, it remains unclear whether its function in the endothelium or the endocardium is required to maintain blood circulation and prevent DA collapse. Our previous work showed that *fli1*-driven *she* expression, which includes vascular endothelium and endocardium, fully rescues circulatory defects in *she* mutants [28]. To determine whether endocardial expression alone was sufficient, we generated *Tg(endo83:she-2A-mCherry)*, which drives *she* expression selectively in endocardial cells (Supplementary Fig. 8). Among 125 transgene-positive embryos obtained from the incross of *she+/-;endo83-she-2A-mCherry* adults, 35 (28%) developed pericardial edema and loss of circulation. None of the 22 genotyped, phenotypically normal embryos were homozygous mutants. In contrast, 22 of 29 embryos with pericardial edema were *she-/-* mutants. Endocardial-specific expression therefore failed to rescue the phenotype. In combination with *fli1*-mediated rescue, these results support a primary requirement for She in the vascular endothelium rather than the endocardium.

### *she* mutant embryos exhibit enlarged glomerular capillary tufts and abnormal pronephric tubule morphology

Because pronephric dysfunction can contribute to edema and circulatory abnormalities [57, 58], we examined kidney morphology at 4 dpf using the *Tg(fli1:lifeact-GFP)* reporter line. The glomerular capillary tuft (GCT) was significantly enlarged in *she* mutant embryos compared with wild-type siblings (Supplementary Fig. 9A, B, E). In addition, the 3G8-labeled proximal pronephric tubules (PT) remained aberrantly straight or uncoiled in *she* mutant embryos, compared to the tightly packed spiral convolutions in wild-type controls (Supplementary Fig. 9C, D). Together, these data indicate that *she* is required for normal proximal tubule morphogenesis and for regulating GCT size. However, whether these defects in pronephric development contribute to DA collapse remains unclear and will require further investigation.

## Discussion

Our findings demonstrate a key role for She in maintaining DA stability and suggest that *she* promotes the accumulation of ECM-producing perivascular fibroblasts, which provide critical support for the DA structure. Before overt circulatory failure, the mutant DA shows enlargement and increased pulsatile diameter change despite no detectable changes in endothelial cell geometry and vessel shape. These abnormalities are accompanied by fewer perivascular stromal cells and reduced perivascular fibrillar collagen gene expression. Partial ablation of mural cells largely preserves DA caliber and circulation, while SNP provides partial rescue. Together, these observations support a model in which inadequate perivascular support makes the developing artery vulnerable to mural cell-associated narrowing (Figure 10). The proposed sequence from fibroblast reduction to altered matrix mechanics and collapse remains to be established directly.

**Figure 10.**
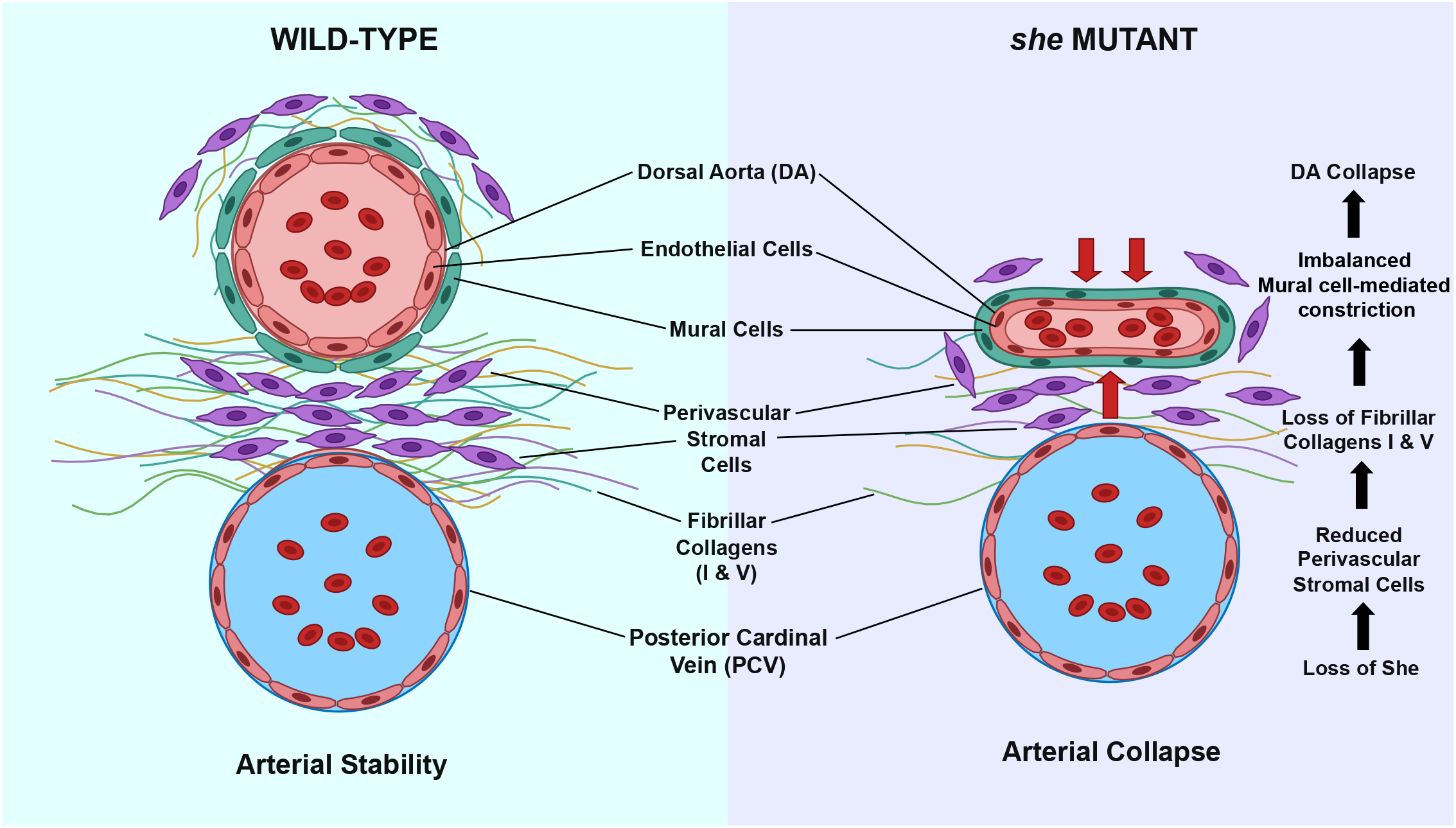
A proposed model showing how She maintains perivascular stromal cells and fibrillar collagens to prevent mural cell-mediated arterial collapse. (Left) Endothelial She expression sustains perivascular stromal cells (purple) in the intervascular space and dorsal to the DA. These stromal cells deposit a network of fibrillar Collagens I and V (wavy multi-colored fibers), which help to maintain arterial stability and luminal patency. (Right) Loss of *she* leads to the depletion of perivascular stromal cells and a corresponding reduction in fibrillar collagens. Without perivascular structural support, normal contractile tone from mural cells contributes to the DA collapse.

The reduction in perivascular stromal cells suggests that She helps coordinate endothelial interactions with the surrounding mesenchyme. Its reported endothelial expression [28, 41] is consistent with a non-cell-autonomous influence on fibroblasts. During normal vascular maturation, the nascent endothelium actively recruits, sustains, and directs the differentiation of surrounding mesenchymal cells through paracrine signaling networks, most notably via PDGF, TGF-β, and Jagged/Notch pathways [21, 59], as well as through biomechanical cues transmitted across the shared ECM [47, 48, 50]. As an SH2 domain-containing adaptor protein, She likely couples endothelial receptor tyrosine kinases to downstream transcriptional or secretory machinery [26, 28, 60, 61]. In the absence of *she*, impaired intracellular signaling within the endothelium may disrupt the synthesis or secretion of essential paracrine trophic factors required for perivascular fibroblast survival, proliferation, or migration [47, 62].

Altered fibrillar collagen expression provides a plausible link between reduced fibroblast abundance and altered vessel behavior. Fibrillar collagens are the primary load-bearing proteins within the vascular ECM [18, 48]. They provide tensile support and limit excessive expansion of vascular tissues [18, 63–65]. Smaller *col1a1* and *col5a1* expression domains, along with lower *col1a1* expression and a reduced Collagen I pro-peptide amount, are therefore consistent with diminished matrix-producing capacity around the mutant DA. However, these measurements do not establish the organization or crosslinking of deposited collagen. Similarly, greater cyclic diameter change is consistent with reduced wall resistance but may also reflect differences in pulse pressure. Direct matrix imaging and pressure-diameter measurements will be needed to determine how the observed cellular changes affect arterial mechanics.

Single-cell RNA sequencing provided complementary evidence for a reduced collagen-rich fibroblast subpopulation. The absence of significant abundance changes among the 27 broad clusters does not conflict with a difference detected after fibroblast subclustering, because a restricted population may be concealed within a larger cluster. Although our previous endothelial rescue experiments demonstrated that She function is required within vascular endothelial cells, we did not identify significant transcriptional changes in these cells. It is plausible that She acts primarily through post-transcriptional mechanisms.

The ablation result provides the strongest functional evidence that mural cells contribute to circulatory failure. Mural cells normally stabilize the developing zebrafish DA and themselves promote collagen deposition [14], yet reducing mural cell coverage preserved the enlarged mutant lumen and largely prevented loss of circulation. One explanation is that mural cell-generated tension narrows a vessel with insufficient structural support. Partial rescue by SNP is consistent with this interpretation. Although *pdgfrb* is expressed in both mural and perivascular fibroblast cell populations, it is expressed at a much higher level in mural cells. Therefore, MTZ treatment of *pdgfrb:Gal4; UAS:mCherry-NTR* embryos might not be expected to cause any significant cell death in the perivascular stromal cell population. In addition, the treatment doses used (50-100 µM) are 100-200-fold lower than the full-strength 10 mM dose; thus, the low treatment doses resulted in only partial ablation of cells with the strongest *pdgfrb+* expression, i.e., mural cells. Thus, our results support the model that mural cells, and likely mural cell-dependent tone, are required for efficient collapse of the *she* mutant DA.

The DA and PCV are located in close proximity at 1-2 dpf stage, but the distance between them substantially increases between 3 and 5 dpf [14, 25]. However, to the best of our knowledge, the cause of this increase has not been previously investigated. Our results show that perivascular stromal cells accumulate between the DA and PCV at this stage. Many of these cells are weakly positive for *pdgfrb:GFP* and collagen expression and thus likely represent a population of perivascular fibroblasts (PVF).

*she* mutants display cardiac defects, including cardiac chamber dilation and atrioventricular regurgitation. However, our results suggest that these cardiac defects are secondary and may be related to failed circulation. Endocardial-specific *she* expression failed to restore circulation, whereas *fli1*-driven expression in the vascular endothelium and endocardium rescues the phenotype [28]. Together, these observations suggest that endocardial expression of *she* alone is insufficient. Cardiac chamber dilation, atrioventricular regurgitation, and pronephric abnormalities could develop in response to altered systemic flow or contribute to its deterioration. It is also possible that *she* function is required within both the endocardium and vascular endothelium. An enhancer driving *she* expression in the vascular endothelium but not the endocardium would be required to answer this question.

The enlarged bulbus arteriosus and increased Elnb expression suggest an additional requirement for She function within the BA. Greater cyclic deformation of the outflow tract may reflect altered loading, changes in matrix properties, or both. Elnb knockdown failed to rescue DA caliber or blood circulation. This suggests that an increase in Elnb is insufficient to explain the circulatory failure, although it could contribute to the local outflow tract phenotype.

In summary, our findings link She function to perivascular fibroblast abundance and the maintenance of arterial patency during development. They support a model in which reduced perivascular support increases susceptibility to mural cell-associated narrowing and circulatory failure (Figure 10). This expands the biological role of She beyond early lumen-size control and identifies endothelial–perivascular interactions as a potential determinant of arterial stability. Since we previously demonstrated that SHE function in restricting the tube size is conserved in human endothelial cell culture, it is likely that SHE may also regulate stromal cell abundance and DA stability in mammalian embryos, although this was not directly tested in the current study.

## Acknowledgements

We thank Dr. Zeenat Diwan and Dr. Arndt Siekmann, University of Pennsylvania for assistance with EC size and shape measurements. We also thank Dr. Geoffrey Burns from Boston Children’s Hospital and Dr. Joshua Waxman from Cincinnati Children’s Hospital for providing the anti-Elnb antibody.

---

**Supplementary Figure 1.**
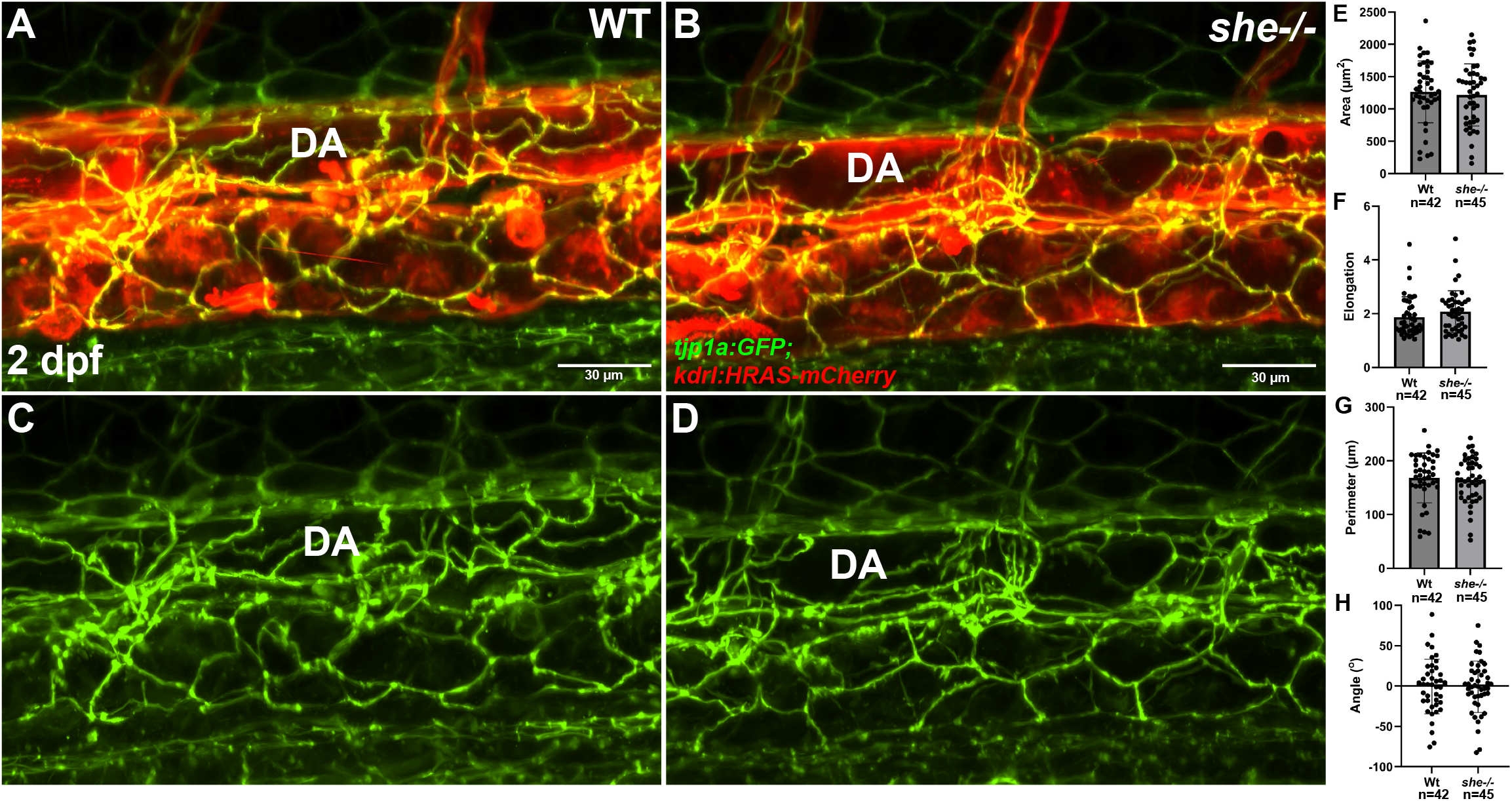
Endothelial cell morphology and alignment in the DA are unaffected in *she* mutant embryos. (A–D) Confocal projections of the DA in wild-type (A, C) and *she* homozygous mutant (B, D) zebrafish embryos at 2 days post-fertilization (2 dpf). (A, B) Merged images of *Tg(tjp1a:eGFP)* (green, labeling tight junctions at cell boundaries) and *Tg(kdrl:HRAS-mCherry)* (red, labeling endothelial cell membranes) in wild-type (A) and *she-/-* (B) embryos. (C, D) Single-channel views of *tjp1a:eGFP* expression alone, illustrating endothelial cell outlines in the DA of WT (C) and *she–/–* (D) embryos. (E–H) Quantification of endothelial cell morphological parameters in the DA of wild-type and *she-/-* embryos, including cell area (E), elongation (F), perimeter (G), and orientation angle (H). Scale bar: 30 µm. n_wt_ = 42 cells from 11 embryos; n*_she-/-_* = 45 cells from 11 embryos; p > 0.05; Student’s t-test. Data were combined from three independent experiments.

**Supplementary Figure 2.**
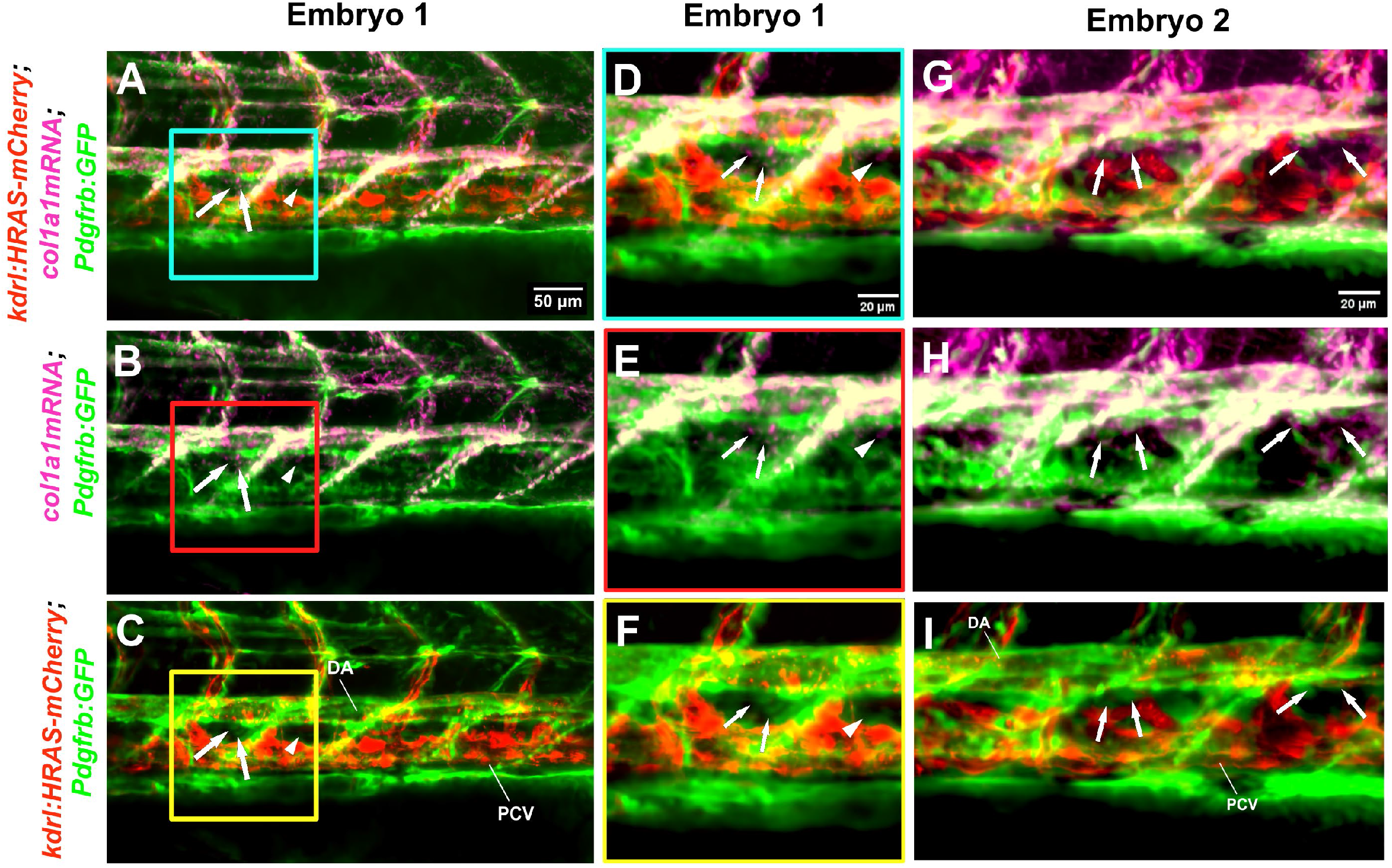
*col1a1* and *pdgfrb:GFP* expression analysis in the stromal cells between the DA and PCV. (A–I) Confocal images of the trunk vasculature in 3 dpf wild-type *Tg(kdrl:HRAS-mCherry; pdgfrb:GFP)* embryos stained for *col1a1* mRNA (magenta) by HCR. (A–C) Overview of the trunk region; boxed areas (cyan, red, yellow) are magnified in D–F. (G–I) High-magnification views of the area between the DA and PCV from an additional embryo. Panels show merged images of all three channels (A, D, G), *col1a1* mRNA and *pdgfrb:GFP* (B, E, H), or *kdrl:HRAS-mCherry* and *pdgfrb:GFP* (C, F, I). Arrows indicate perivascular stromal cells between the DA and PCV coexpressing *col1a1* mRNA and *pdgfrb:GFP*. Arrowheads (A–F) mark a *col1a1* mRNA-positive cell lacking visible *pdgfrb:GFP* expression. Scale bar: 50 µm (A-C) and 20 µm (D-I).

**Supplementary Figure 3.**
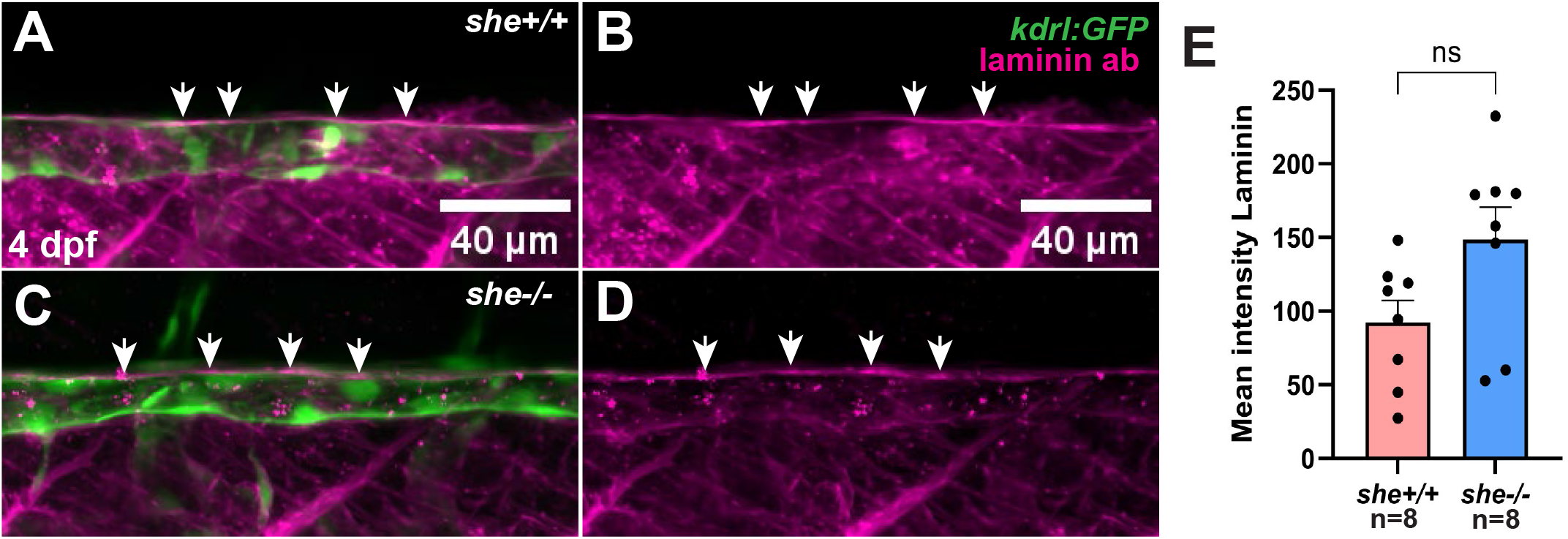
Laminin deposition around the DA is not significantly affected in *she* mutant embryos. (A–D) Representative confocal projections of the trunk vasculature in *she+/+* sibling (A, B) and homozygous *she-/-* mutant (C, D) zebrafish embryos at 4 dpf immunostained with an anti-laminin antibody. Vascular endothelium is labeled by *kdrl:GFP* (green), while laminin immunostaining is in magenta (Alexa Fluor 647). A and C show merged *kdrl:GFP* and laminin staining images, while B and D show single-channel laminin staining images in *she+/+* and *she-/-* embryos, respectively. Arrows point toward laminin staining around the DA. (E) Quantification of the mean fluorescence intensity of laminin staining around the dorsal aorta in *she+/+* (n = 8) and *she-/-* (n = 8) embryos at 4 dpf. Scale bar: 40 µm. ns, not significant; Student’s t-test.

**Supplementary Figure 4.**
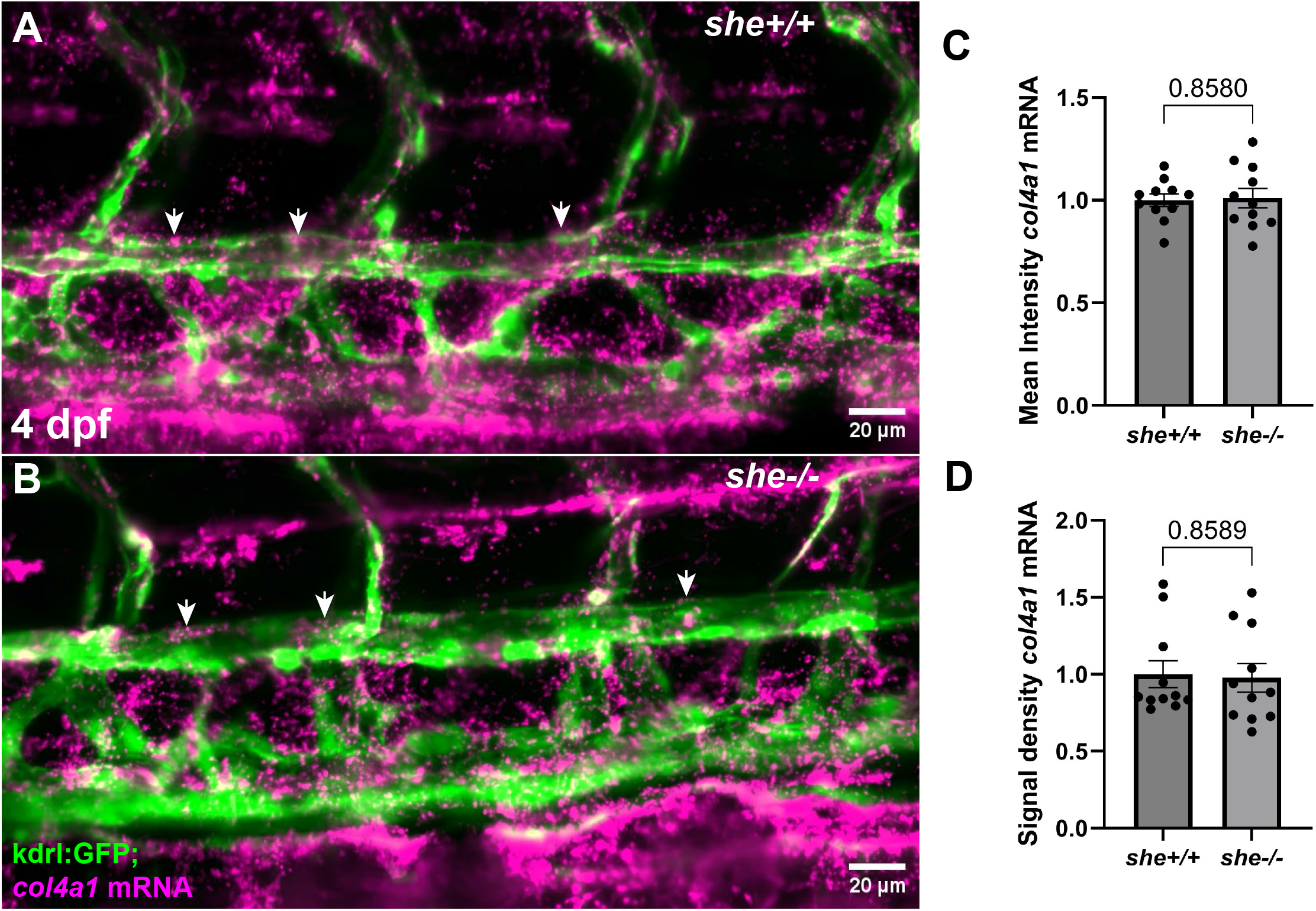
Expression of the basement membrane component *col4a1* around the dorsal aorta is unaffected in *she* mutant embryos. (A, B) Representative confocal images of FISH for *col4a1* mRNA in sibling *she+/+* (A) and homozygous *she-/-* mutant (B) embryos expressing the endothelial reporter *Tg(kdrl:GFP)* at 4 dpf, showing *col4a1* transcript distribution around the dorsal aorta (white arrows). (C, D) Quantification of normalized mean fluorescence intensity of *col4a1* mRNA (C) and *col4a1* mRNA signal density (D) around the DA in *she+/+* (n = 11) and *she-/-* (n = 11) embryos at 4 dpf. Scale bar: 20 µm. p-values were determined by Student’s t-test.

**Supplementary Figure 5.**
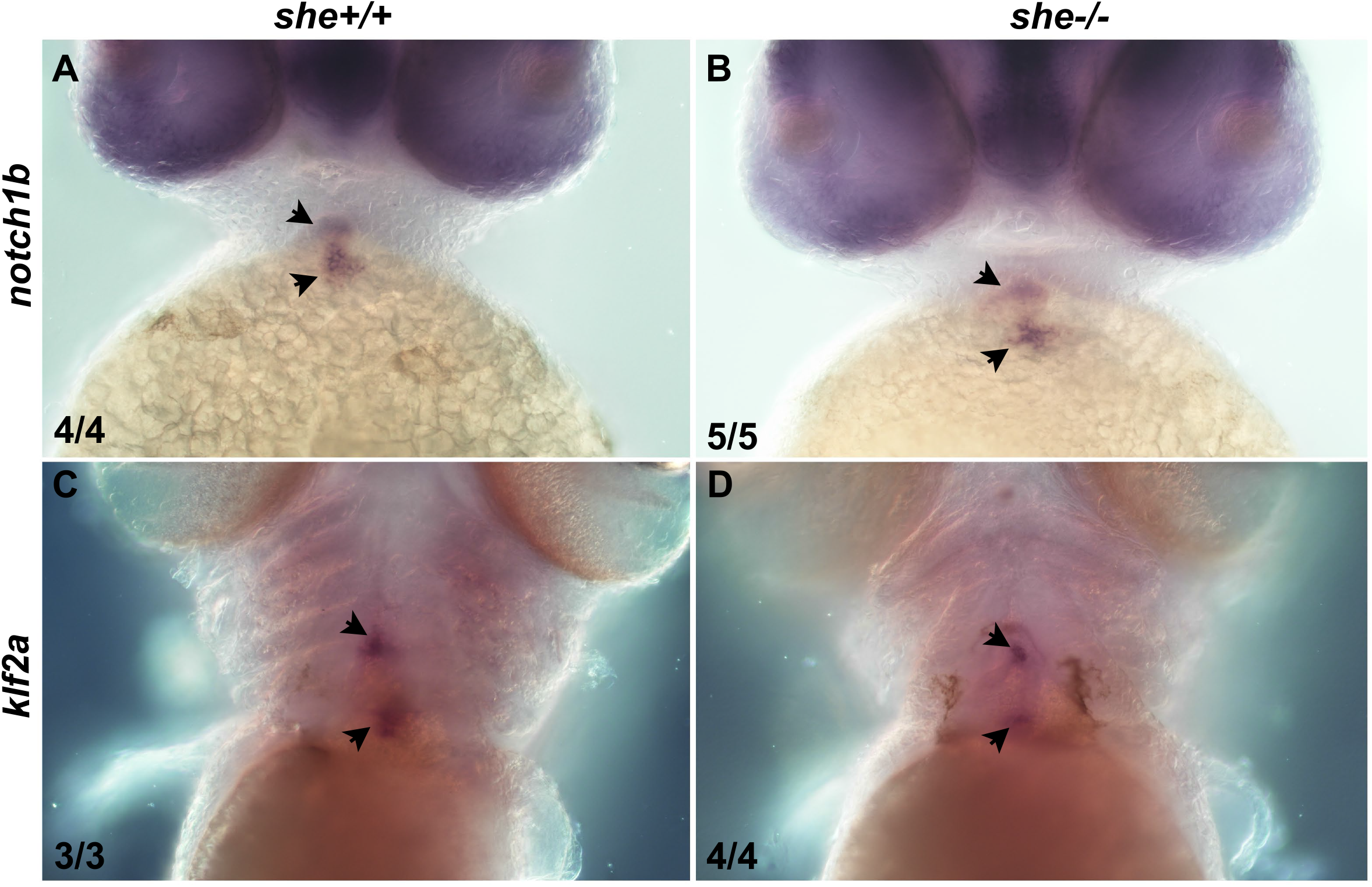
Expression of the cardiac valve-associated markers *notch1b* and *klf2a* is unaffected in *she* mutant embryos. (A–D) Representative ventral views of the cardiac region following whole-mount in situ hybridization (WISH) for *notch1b* (A, B) and *klf2a* (C, D) in wild-type sibling (A, C) and homozygous *she* mutant (B, D) zebrafish embryos at 2 dpf (A, B) and 4 dpf (C, D). Embryos were obtained from an in-cross of heterozygous *she+/-* parents and genotyped afterward. Black arrowheads point to localized transcript expression in the endocardial cushions of the outflow tract (upper arrowhead) and atrioventricular canal (lower arrowhead). Both *notch1b* and *klf2a* show comparable expression levels and spatial restriction between *she+/+* and s*he-/-* embryos. The number of embryos exhibiting the representative genotype and phenotype out of the total number examined is shown in the lower left corner.

**Supplementary Figure 6.**
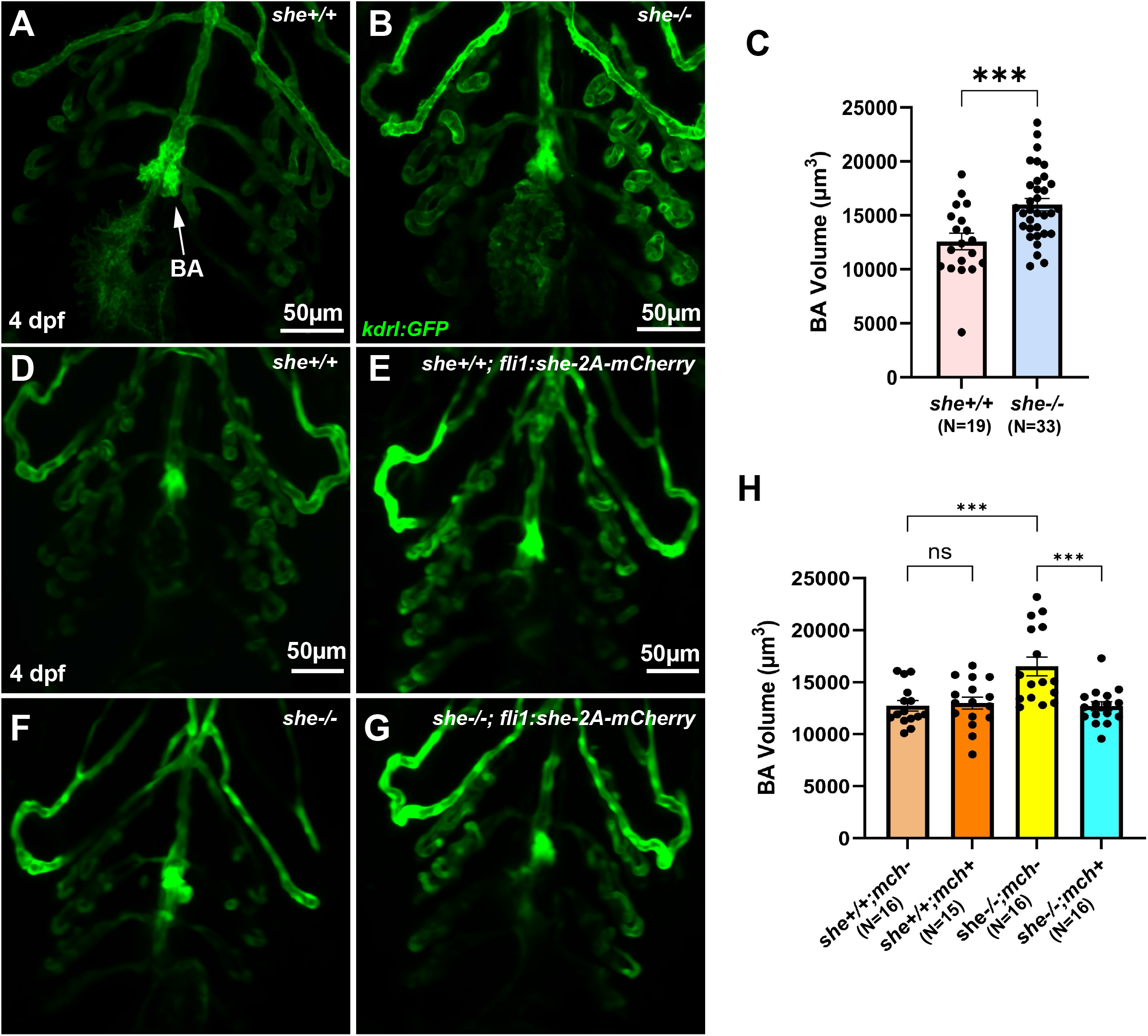
*she* mutants exhibit enlargement of the bulbus arteriosus (BA). (A-C) Quantitative analysis of BA size using Imaris-based 3D volumetric quantification of representative confocal images from wild-type (A) and *she* mutant (B) embryos in a *kdrl:GFP* background at 4 dpf. n*_she+/+_* =19; n*_she-/-_* =33; ***p<0.001, Student’s t-test. Data were combined from two independent experiments. (D-H) Representative confocal images (D-G) and quantitative analysis (H) of BA volume in wild-type (D, E) and *she* mutant (F, G) embryos with (E, G) and without (D, F) expression of the rescue transgene *fli1:she-2A-mCherry* in the *kdrl:GFP* reporter background at 4 dpf. *fli1:she-2A-mCherry* expression did not alter BA dimensions in wild-type embryos, but successfully restored BA size in *she* homozygous mutants (H). Scale bar: 50 µm. n*_she+/+;mch_*-=16, n*_she+/+;mch+_*=15, n*_she-/-;mch-_*=16, n*_she-/-;mch+_*=16; ns: not significant; ***p<0.001; one-way ANOVA followed by Šidák’s multiple comparisons test. Data were combined from two independent experiments.

**Supplementary Figure 7.**
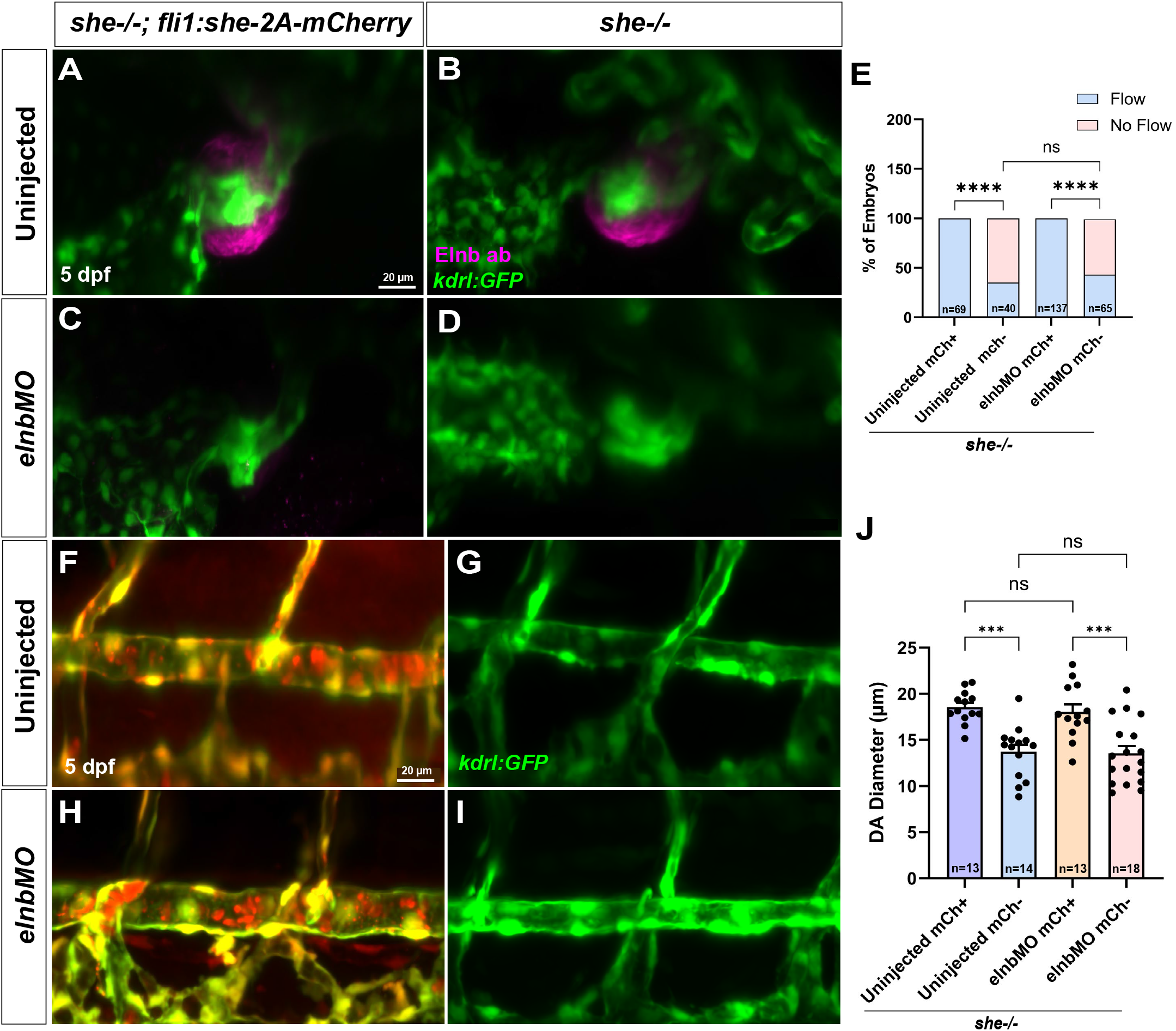
Morpholino knockdown of *elnb* does not rescue DA collapse and circulatory failure in *she* mutant embryos. (A–D) Representative confocal images of the BA in 5 dpf uninjected (A, B) and *elnb* morpholino-injected (*elnb*MO; C, D) *she-/-; Tg(kdrl:GFP)* zebrafish embryos carrying (A, C; *she-/-; fli1:she-2A-mCherry*, rescued controls) or lacking (B, D; *she-/-*, non-rescued mutants) the endothelial rescue transgene. The endothelium and endocardium are labeled by *kdrl:GFP* (green), and whole-mount immunofluorescence detects Elnb protein (magenta). Injection of *elnb*MO markedly depletes Elnb immunoreactivity in the BA (C, D) compared to uninjected siblings (A, B). (E) Quantification of the percentage of 5 dpf *she-/-* embryos with or without blood circulation across four groups: uninjected mCherry-positive (n=69), uninjected mCherry-negative (n=40), *elnb*MO-injected mCherry-positive (n=137), and *elnb*MO-injected mCherry-negative (n=65). ns, not significant; ****p < 0.0001, Fisher’s exact test. (F–I) Representative confocal images of the DA of 5 dpf uninjected (F, G) and *elnb*MO-injected (H, I) *she-/-*; Tg(kdrl:GFP) embryos carrying (F, H) or lacking (G, I) the *fli1:she-2A-mCherry* rescue transgene. (J) Quantification of DA luminal diameter at 5 dpf in uninjected mCherry (mCh)+ (n = 13), uninjected mCh- (n = 14), *elnb*MO mCh+ (n = 13), and *elnb*MO mCh- (n = 18) *she-/-* embryos. Scale bar: 20 µm. ns, not significant; ***p < 0.001; one-way ANOVA followed by Šidák’s multiple comparisons test.

**Supplementary Figure 8.**
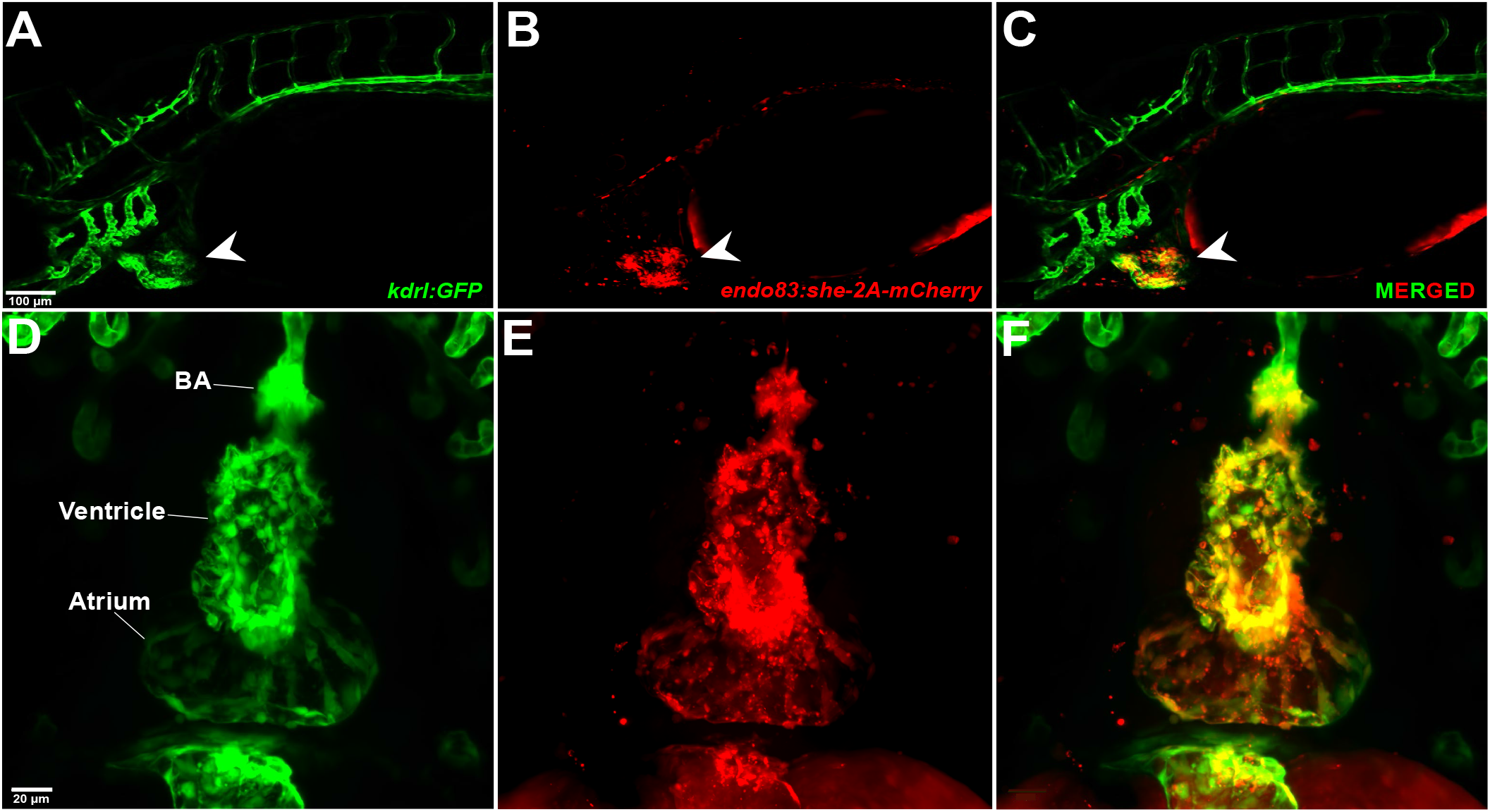
Expression analysis of the endocardial-specific *Tg(endo83:she-2A-mCherry)* transgenic zebrafish line. (A–C) Representative lateral confocal projections of a *Tg(kdrl:GFP; endo83:she-2A-mCherry)* zebrafish embryo at 4 dpf. (A) *Tg(kdrl:GFP)* expression (green) labels the vascular endothelium and cardiac endocardium. (B) *Tg(endo83:she-2A-mCherry)* expression (red) is driven by the endocardial-specific *endo83* enhancer coupled to a minimal basic promoter. (C) Merged image demonstrating that within the cardiovascular system, *endo83:she-2A-mCherry* expression colocalizes with *kdrl:GFP* in the heart (white arrowheads in A–C) and is absent from the trunk vasculature. Note the strong nonvascular red fluorescence in the yolk extension. (D–F) High-magnification ventral confocal projections of the heart in a *Tg(kdrl:GFP; endo83:she-2A-mCherry)* embryo at 4 dpf showing *kdrl:GFP* (green; D), *endo83:she-2A-mCherry* (red; E), and the merged channels (F). *endo83:she-2A-mCherry* is robustly expressed throughout the endocardium of the atrium and ventricle, as well as the bulbus arteriosus (BA), while being absent from the surrounding pharyngeal arch vasculature. Scale bar: 20 µm.

**Supplementary Figure 9.**
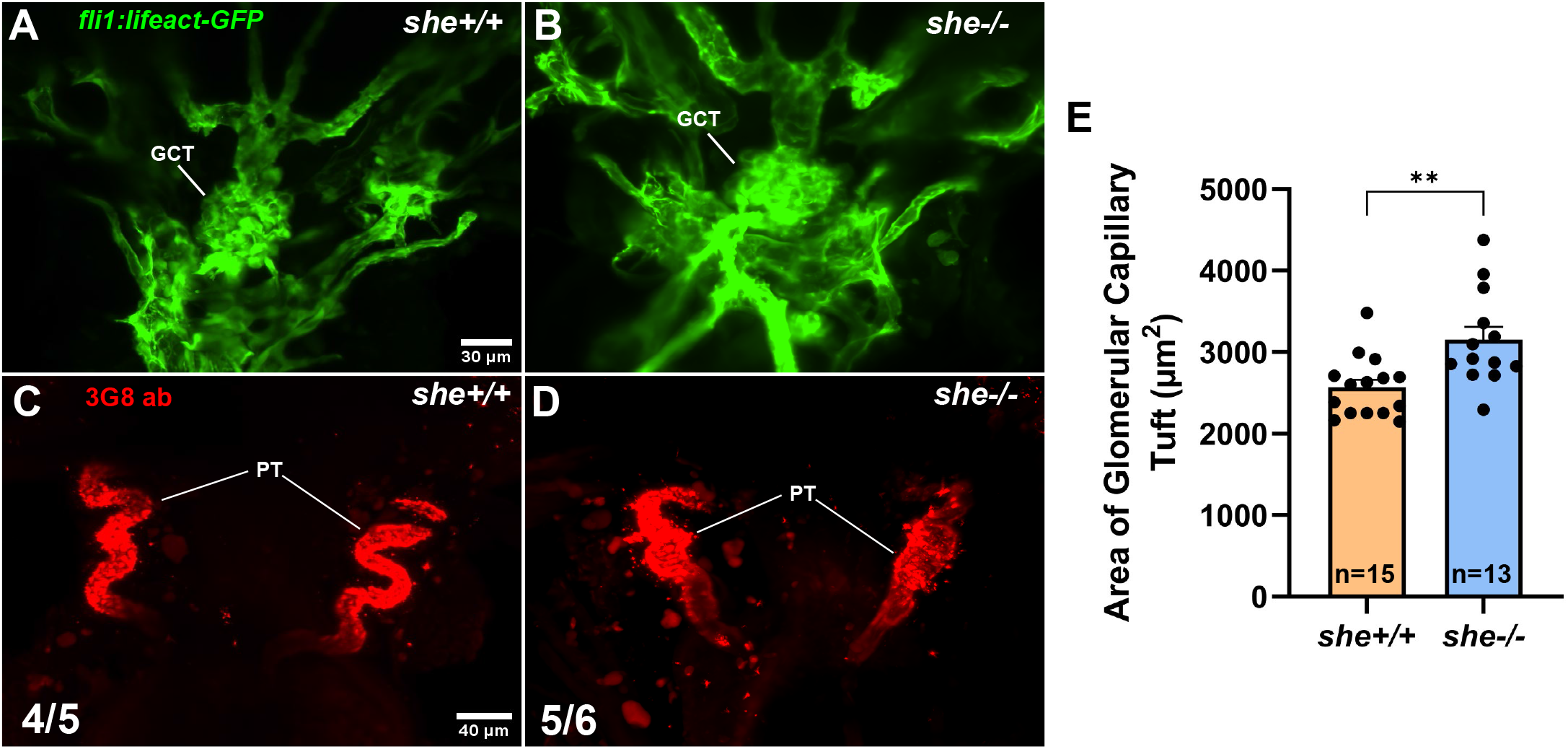
Loss of *she* results in glomerular capillary tuft enlargement and impaired proximal pronephric tubule convolution. (A, B) Representative confocal projections of the glomerular capillary tuft (GCT) in *she+/+* sibling (A) and homozygous *she-/-* mutant (B) zebrafish embryos expressing the *Tg(fli1:lifeact-GFP)* endothelial reporter at 4 days post-fertilization (4 dpf). (C, D) Whole-mount immunofluorescence staining with the 3G8 monoclonal antibody (red) labeling the proximal pronephric tubules (PT) in *she+/+* (C) and *she-/-* (D) embryos at 4 dpf. While *she+/+* embryos exhibit normally convoluted proximal tubules (4/5 embryos; C), *she-/-* mutants display aberrantly straight, uncoiled tubules (5/6 embryos; D). The number of embryos exhibiting the representative phenotype out of the total number examined is shown in the bottom left corner. (E) Quantification of the glomerular capillary tuft area (µm²) in *she+/+* (n = 15) and *she-/-* (n = 13) embryos at 4 dpf. Scale bar: 30 µm (A, B) and 40 µm (C, D). **p < 0.01, Student’s t-test. Data were combined from two independent experiments.

## Movie Legends

**Supplementary Movie 1. Time-lapse imaging of progressive DA collapse in *she* mutant zebrafish embryos from 3 to 5 dpf.**

Representative confocal time-lapse movie of the trunk vasculature in a rescued *she-/-; fli1:she-2A-mCherry* embryo (top panel) and a non-rescued homozygous *she-/-* mutant sibling (bottom panel) in a *Tg(kdrl:GFP)* vascular endothelial reporter background imaged from 3 to 5 dpf. In the rescued *she-/-; fli1:she* control embryo, the DA maintains a patent lumen throughout the 3–5 dpf imaging period. In contrast, the DA in the *she-/-* mutant embryo initially shows an open lumen at 3 dpf but progressively narrows and ultimately collapses into a thin, constricted cord by 5 dpf. Images were acquired every 30 minutes over 40 hours and are played back at 10 frames per second.

**Supplementary Movie 2. Live imaging capturing pulsatile distensibility of the DA in zebrafish embryos.** Representative real-time fluorescence movie of the trunk vasculature in a wild-type sibling (left) and a homozygous *she-/-* mutant (right) zebrafish embryo in a *Tg(kdrl:GFP)* vascular endothelial reporter background at 3.5 dpf. The DA and PCV are labeled. Images were captured every 68 milliseconds over 10.5 seconds and are played back at 7 frames per second.

**Supplementary Movie 3. Live brightfield imaging capturing heartbeats for distensibility analysis of the atrium, ventricle, and bulbus arteriosus in zebrafish embryos.**

Representative high-speed brightfield time-lapse movie (ventral view) of the beating heart in a wild-type sibling (left) and a homozygous *she-/-* mutant (right) zebrafish embryo at 4 dpf. The atrium, ventricle, and bulbus arteriosus (BA) are labeled. Note the pronounced distension of the atrium and BA as blood is pumped through the cardiac chambers in the *she-/-* embryo. Images were captured every 62 milliseconds over 10 seconds and are played back at 10 frames per second.

**Supplementary Movie 4. High-speed brightfield imaging reveals retrograde blood flow across the atrioventricular valve in *she-/-* mutant zebrafish embryos.**

Representative high-speed brightfield time-lapse movie of the beating heart in a wild-type sibling (left) and a homozygous *she-/-* mutant (right) zebrafish embryo at 4 dpf. The atrium, ventricle, and bulbus arteriosus (BA) are indicated. While the wild-type sibling exhibits unidirectional blood flow from the atrium into the ventricle and out through the BA during each cardiac cycle, the *she-/-* mutant embryo displays pronounced retrograde blood flow (backflow, indicated by the white arrow) from the ventricle back into the atrium across the atrioventricular canal during ventricular systole. Images were captured every 6.4 milliseconds over 3 seconds and are played back at 20 frames per second.

